# Unravelled proteins form blobs during translocation across nanopores

**DOI:** 10.1101/2024.01.23.576815

**Authors:** Adina Sauciuc, Jacob Whittaker, Matthijs Tadema, Kasia Tych, Albert Guskov, Giovanni Maglia

## Abstract

The electroosmotic-driven transport of unravelled proteins across nanopores is an important biological process that is now under investigation for the rapid analysis and sequencing of proteins. For this approach to work, however, it is crucial that the polymer is threaded in single file. Here we found that, contrary to the electrophoretic transport of charged polymers such as DNA, during polypeptide translocation blob-like structures typically form inside nanopores. Comparisons between different nanopore sizes, shapes and surface chemistries showed that under electroosmotic-dominated regimes single-file transport of polypeptides can be achieved using nanopores that simultaneously have an entry and an internal diameter that is smaller than the persistence length of the polymer, have a uniform non-sticky (*i*.*e*. non-aromatic) nanopore inner surface, and using moderate translocation velocities.

## Introduction

Polymer transport across nanopores has been extensively studied using uniformly charged polymers such as DNA. The electrophoretic translocation of single-stranded DNA (ssDNA) has been reported for a range of nanopores including alpha hemolysin (αHL)^1^, MspA^2^ and FraC^3^. Despite these nanopores have different entry diameters (of 2 nm, 4 nm and 6 nm, respectively) and constriction size (1.6 nm, 1 nm and 1.2 nm, respectively), electrophoretic driven ssDNA transport has been always reported as single file in all nanopores.
Proteins, however, do not have a uniform charge, and their transport across a nanopore can only be obtained using electroosmotic forces^4–8^. Exploring the electroosmotically-driven transport of unfolded proteins across CytK, MspA, aerolysin and lysenin nanopores, to our surprise, we found that polypeptides typically form blob-like coiled structures inside the nanopore during translocation. Single-file transport of unfolded proteins, which is crucial in protein sequencing applications, is only observed using a nanopore with a specific size, shape and chemical composition, and under moderate translocation speeds.

## Results

### CytK structure and protein transport

It was recently shown that polypeptide translocation can be observed using CytK-2E-4D (shortened as CytK-4D)^7^ where 4 Asp residues were introduced in the β-barrel region of the nanopore to introduce a strong electro-osmotic flow (EOF) [P(K^+^)/P(Cl^-^) ratio of 4.0 ± 0.1]. The latter induced the translocation of unstructured protein substrates even against relatively strong electrophoretic forces (EPF). Similar results were observed using WT-αHL nanopores, a homologue of CytK, during translocation induced by both EP and EPF forces^4^.

To identify the characteristics in nanopores that allow for single-file translocation, we resolved the structure of CytK-K128D-T147D-K155D, a CytK mutant with enhanced solubility which allows the transport of unfolded proteins. The structure was resolved at a resolution of 4.4 Å (**Table S1**) and, as predicted by homology modelling ^7,9^, the nanopore was heptameric and very similar to α-hemolysin (**Figure 1A, S1**).

**Figure 1.**
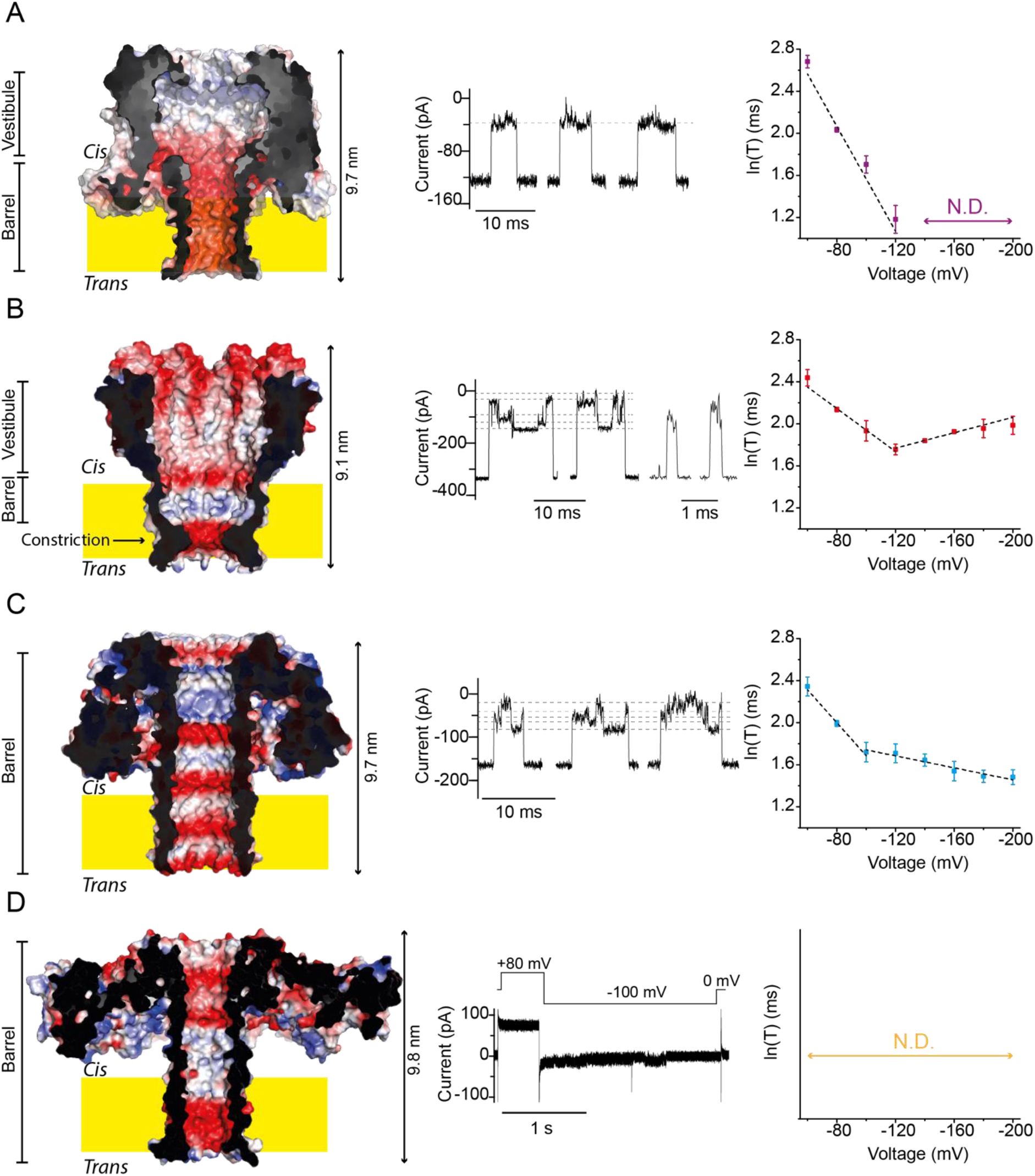
Translocation of the malE219a protein across highly cation selective nanopores: CytK-4D (**A**), MspA WT (**B**), lysenin WT (**C**) and aerolysin 1D-3N (**D**), coloured by calculated electrostatic potential. The middle traces show representative events obtained upon addition of malE219a on the cis side of the nanopores. The right graphs indicate the dwell time (τ) dependence on the applied potential corresponding to the translocation through the nanopores. Data were collected in 1 M KCl, 2 M urea, 15 mM HEPES buffer pH 7.5. Traces were sampled at 50 kHz and filtered at 10 kHz.

### Protein translocation across CytK, MspA, lysenin and aerolysin nanopores

In this work we compared the translocation of malE219a under protein unfolding conditions (2 M urea, 1 M KCl and 15 mM HEPES buffer pH 7.5)^7^ using four different nanopores: CytK-4D (**Figure 1A**), MspA (**Figure 1B**), lysenin (**Figure 1C**) and an engineered aerolysin nanopore (**Figure 1D**). The CytK-4D nanopore comprises of a narrow entry of 1.7 nm followed by a vestibule of ∼3 nm at its widest point and a ∼1.5 nm β-barrel (1.2 nm at its narrowest point). MspA-M2 mutant,^2^ which has been employed in DNA^10^ and peptide^11–14^, has an asymmetric hour-glass shape and comprises a broad vestibule (∼4 nm at the *cis* entry), followed by a ∼3 nm β-barrel region and a 1.2 nm constriction at the *trans* exit (**Figure 1B**). To test for electroosmotically-driven transport of polypeptides, here we employed the highly cation selective [P(K^+^)/P(Cl^-^) of 4.1 ± 0.2] wild-type nanopore. The lysenin nanopore has a cylindrical shape, where the top half contains two 1.5 nm constrictions defined by K37 and K45, while the bottom half of the nanopore is ∼2 nm wide (**Figure 1C**). Lysenin is cation selective, as indicated by the reported P(K^+^)/P(Cl^-^) ratio of 7.5 ± 0.5^15^. Finally, we used the aerolysin nanopore, which has been employed in peptide identification^9^ and amino acid recognition^16^, and has been used to translocate proteins unfolded by guanidinium chloride^5,6^. This nanopore has a 1.2 – 1.5 nm wide cylindrical lumen (defined as the water facing region of the nanopore, **Figure 1D**). Aer-WT, however, is weakly anion selective [P(K^+^)/P(Cl^-^) of 0.76 ± 0.07^9^), and extensive engineering was required to promote a strong EOF for the translocation of unstructured proteins in the absence of denaturant or in urea. R220, K242 and R282 were substituted with Asn and K238 with Asp, which created a highly cation selective nanopore [aer-1D-3N, [P(K^+^)/P(Cl^-^) 3.95 ± 0.08, **Figure 1D**]. The introduction of an additional Asp at position 272 further increased the cation selectivity to 4.68 ± 0.19 (aer-2D-3N).

As also reported earlier^7^, in the -60 to -120 mV range the translocation of unfolded malE219a across CytK-4D induced well-defined blockades with an average excluded current of 71.9 ± 0.8 % at -80 mV. The dwell time associated with the events decreased sharply with the applied potential from 24.4 ± 6.7 ms to 3.3 ± 0.5 ms (**Figure 1A**), demonstrating the translocation of the polypeptide across the nanopore. Long-lasting current events were occasionally observed, which were characterised by the following pattern: an initial full block level followed by a subsequent ∼70 % I_ex%_ level [defined as (I_o_ - I_B_)/ I_o_ percent, with I_o_ the open pore current and I_B_ the blocked pore current] and eventually reaching full block or the open pore current (**Figure S2**). These long-lasting events were observed with other unfolded proteins (**Figure S2**), increased with the applied bias and became prominent at voltages above -120 mV (**Figure S3**), preventing the use of these voltage regimes for polypeptide characterisation.

Unfolded malE219a also induced blockades to WT-MspA nanopore starting at -60 mV. Two types of events occurred in the -60 to -120 mV range, which we term type I and type II events (**Figure 1B, S4**). Type I events were characterised by frequent shifts between levels. The shallowest and the deepest levels were rather well-defined and predominant and the transitions between these levels consisted of various intermediate levels (**Figure S4**). Under this voltage regime the dwell time decreased from 11.5 ± 0.9 ms to 5.8 ± 0.3 ms (**Figure 1B**) which is consistent with protein translocation. Type II events were shorter than type I and associated with a smaller excluded current. Their dwell time increased with the potential (**Figure S4**), suggesting these events may correspond to the capture and release back in the *cis* compartment. At higher applied potentials, only type I events were observed, whose dwell time gradually increased to reach 7.3 ± 0.6 ms at -200 mV. This behaviour is consistent with the unfolded protein not translocating under high voltage regimes. The average I_ex%_ increased with the applied potential across the whole range: from 70.2 ± 1.7 % at -60 mV to 87.2 ± 1.7 % at -200 mV (**Figure S4**).

Unfolded malE219a induced multi-level events to WT-lysenin nanopores throughout the entire range of tested potentials (**Figure 1C**). Events appeared at -60 mV, and their dwell time decreased during the entire range of the applied bias applied (up to -200 mV), suggesting that the protein translocated through the nanopore across the entire voltage range. Nonetheless, two regimes were still observed with different trends in translocation times with a turning point at -100 mV (**Figure 1C, S5**). The dwell time in the first voltage range decreased from 10.5 ± 0.9 ms at -60 mV to 5.6 ± 0.6 ms at -100 mV. After the turning point at -100 mV, the dwell time weakly decreased from 5.5 ± 0.5 ms at -120 mV to 4.4 ± 0.3 ms at -200 mV. The I_ex%_ increased over the whole range of sampled potentials: from 69.5 ± 0.3 % at -60 mV to 73.9 ± 3.1 % at -200 mV.

The addition of the same unfolded protein to aerolysin 1D-3N or 2D-3N nanopores resulted in long full block events at any of the sampled potentials (**Figure 1D, S6**) that most often required reverting the applied potential to obtain the open pore current. This is surprising given that unfolded protein translocation was observed with WT-Aer in the presence of molar concentrations of guanidine-chloride (a charged chaotropic agent). Increasing the salt concentration to 1.8 M KCl led to the appearance of translocation events, characterised by a reduced dwell time as the applied potential increased (622 ± 25.5 ms at -80 mV up to 181 ± 3.9 ms at -140 mV, aerolysin 1D-3N, **Figure S7**), suggesting that there is an electrostatic interaction between the nanopore ∼14 nm cap domain and the unfolded polypeptide during translocation.
The importance of the surface interaction prior to threading was tested by translocating the unfolded protein substrate from the *trans* side, where the polypeptide is expected to interact with the lipid bilayer (**Figure 1**). CytK-4D, MspA, 1D-3N-Aer, 2D-3N Aer and lysenin blockades were either long and often irreversible (i.e., the applied potential had to be reversed to reopen the nanopore), or associated with multiple levels, (**Figure S8**-**S11**) suggesting that the polypeptides likely interacted with the lipid bilayer prior entering the nanopore.

### Stretched and coiled protein transport

Proteins unfolded by denaturant exhibit polymer-like behaviour, whose flexibility can be quantified through their persistence length (L_p_). In polypeptides the L_p_ is influenced by multiple factors such as the amino acid composition ^17,18^, the ionic strength of the solution ^17^, and nanoscale confinement ^5,19,20^, and it has been reported to range from one^17^ to ten^21^ amino acids under applied force. A widely accepted persistence length for unstructured polypeptides in solution is five residues^22^, equivalent to a contour length of 1.7 – 1.8 nm (3.4 Å^23^ – 3.6 Å^24–26^ per amino acid). In dilute conditions (consistent with the 0.3 μM concentration used here), polypeptides are in a predominantly coiled state characterised by a radius of gyration (Rg) ranging from ∼3 nm for a 124 amino acid (AA) protein to ∼8 nm for a 549 AA protein^27^. Interpolation from previous data^27^ suggests that unfolded malE219a (412 amino acids) has a Rg of ∼7 nm (**Figure S12A**).

Under confinement, *i*.*e*., inside a nanopore, If the pore diameter (D) is much smaller than the persistence length (D << L_p_) the polymer follows the Odijk regime, adopting a stretched configuration. For D >> L_p_, the de Gennes regime is observed, where the polymer forms structures referred to as blobs^5,28^ (**Figure S12B**). In intermediate regimes, as in the case considered here of a polypeptide inside a nanopore, non-uniform blobs or back-folding may occur^29^. However, the nanopore shape, the solvent flow and the friction of the polymer during translocation are also expected to influence blob formation^30,31^.

MspA has a near conical shape with a large ∼4 nm *cis* entry and a narrow 1 nm constriction, suggesting that unfolded proteins are likely to enter the nanopore in a partially coiled state (Rg ∼7 nm). The multilevel events can then be explained by the unravelling or translocation of blobs of different sizes (**Figure 2A**). The shallow current blockade not associated with translocation (type II events) most likely reflect coils protruding and then retracting the *cis* side. One the other hand, the electroosmotic flow and electrophoretic forces, can also compact the coil further inside the nanopore lumen driving the formation of blob-like structures (**Figure 2A**). Upon reaching the constriction, the blobs will then unravel or translocate as blobs of different size, in turn explaining multiple-level events. The longer dwell times and increased I_ex%_ at V > -120 mV likely resulted from compression of the blobs against the constriction, which in turn reduce the ionic current and weaken the EOF, hindering translocation under these voltages.

**Figure 2.**
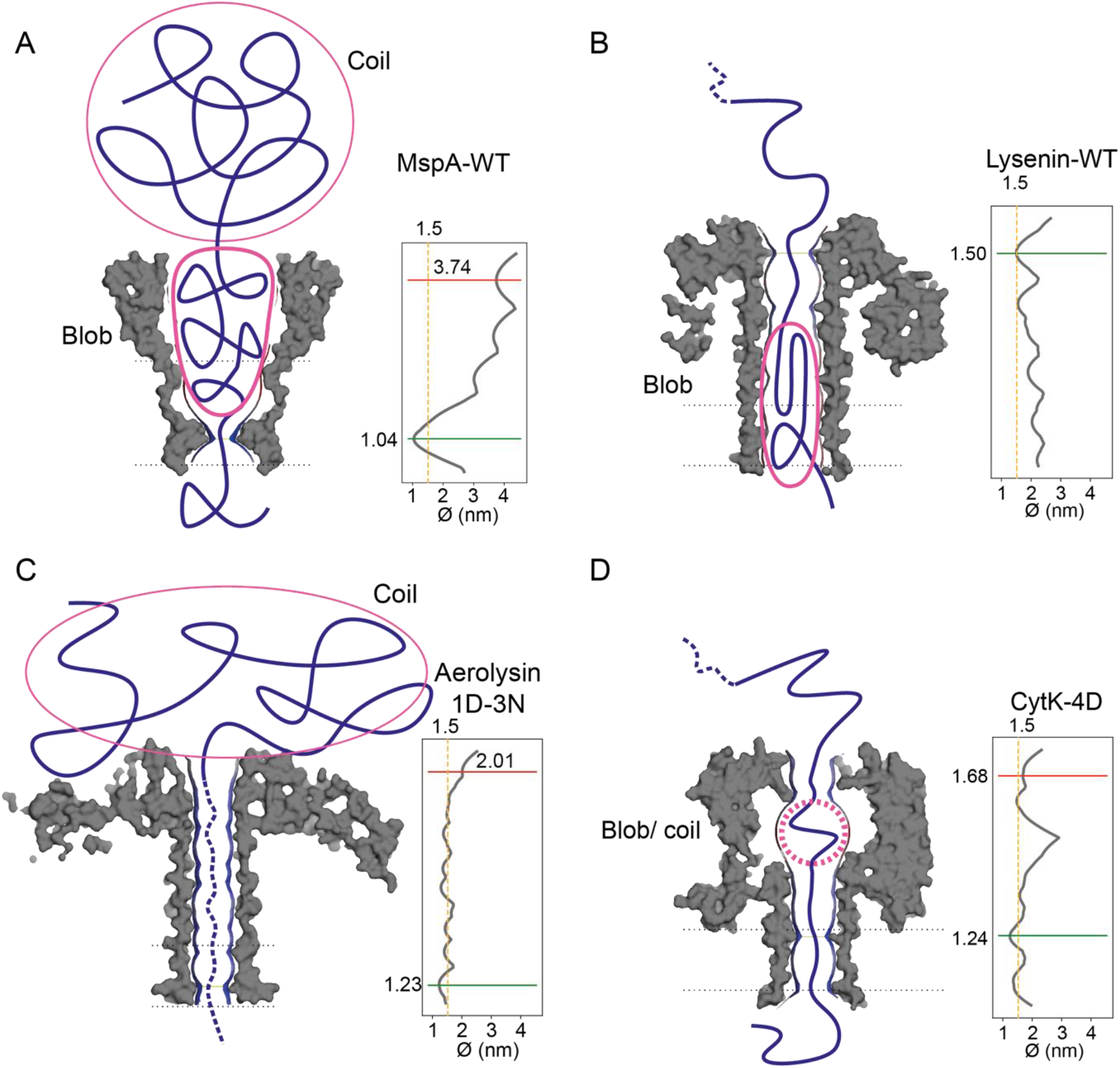
Blob formation during protein translocation across nanopores. In MspA (**A**), the entry of the pore is ∼3-fold larger than the persistance length of a polypeptide, suggesting that blobs can be formed in the lumen of the nanopore. In lysenin (**B**), the nanopore entry is similar to the persistance length of the polypeptide, suggesting that the polymer enters the nanopore as a single file. Then the lysenin barrel opens to about 2 nm and coils can form. In aerolysin 1D-3N (**C**), the polypeptide may coil at the nanopore entry due to the size and composition of the cap domain. In CytK-4D (**D**), the nanopore has two narrow points with diameters smaller than the L_P_ of the polypeptide, possibly preventing the formation of blobs.

Lysenin has a cylindrical shape, whose entry diameter (1.5 nm) is similar to the polymer’s persistence length (∼1.7 nm), suggesting the polypeptide undergoes a certain degree of unfolding before threading^30,31^. Blob-like structures or back folding might then occasionally occur in the second half of the nanopore where an enlargement is observed (**Figure 2B**), explaining the multilevel current events. Polypeptide translocation appears to occur across the entire voltage range tested, suggesting that blob-like structures that may form can be pushed out of the nanopore.

Aerolysin 1D-3N consists of a 1.2-1.5 nm wide barrel, which should be ideal for linear transport. Urea-unfolded proteins, however, can only translocate at high ionic strength. This is likely because of the presence of a large cap domain (14 nm wide) which induces electrostatic interactions with the polypeptide prior translocation (**Figure 2C**).

CytK-4D is consistently ∼1.5 nm wide (**Figure 2D**). In the -60 mV / -120 mV regime, relatively uniform current blockades suggest that blob-like structures are not formed (**Figure 2D, 3**). Above -120 mV, many long-lasting events permanently blocked the nanopore. A likely explanation is that higher transport speed increases the likelihood of sudden changes in the speed of transport, which could favour the formation of local blob-like/coiled structures, possibly in the vestibule of the nanopore (see below). Such structures would then reduce the ionic current and the corresponding EOF preventing their unravelling. This reduction may further hamper the unravelling of coils from solution as they approach the nanopore and prevent the transport across the nanopore (**Figure 3**).

**Figure 3.**
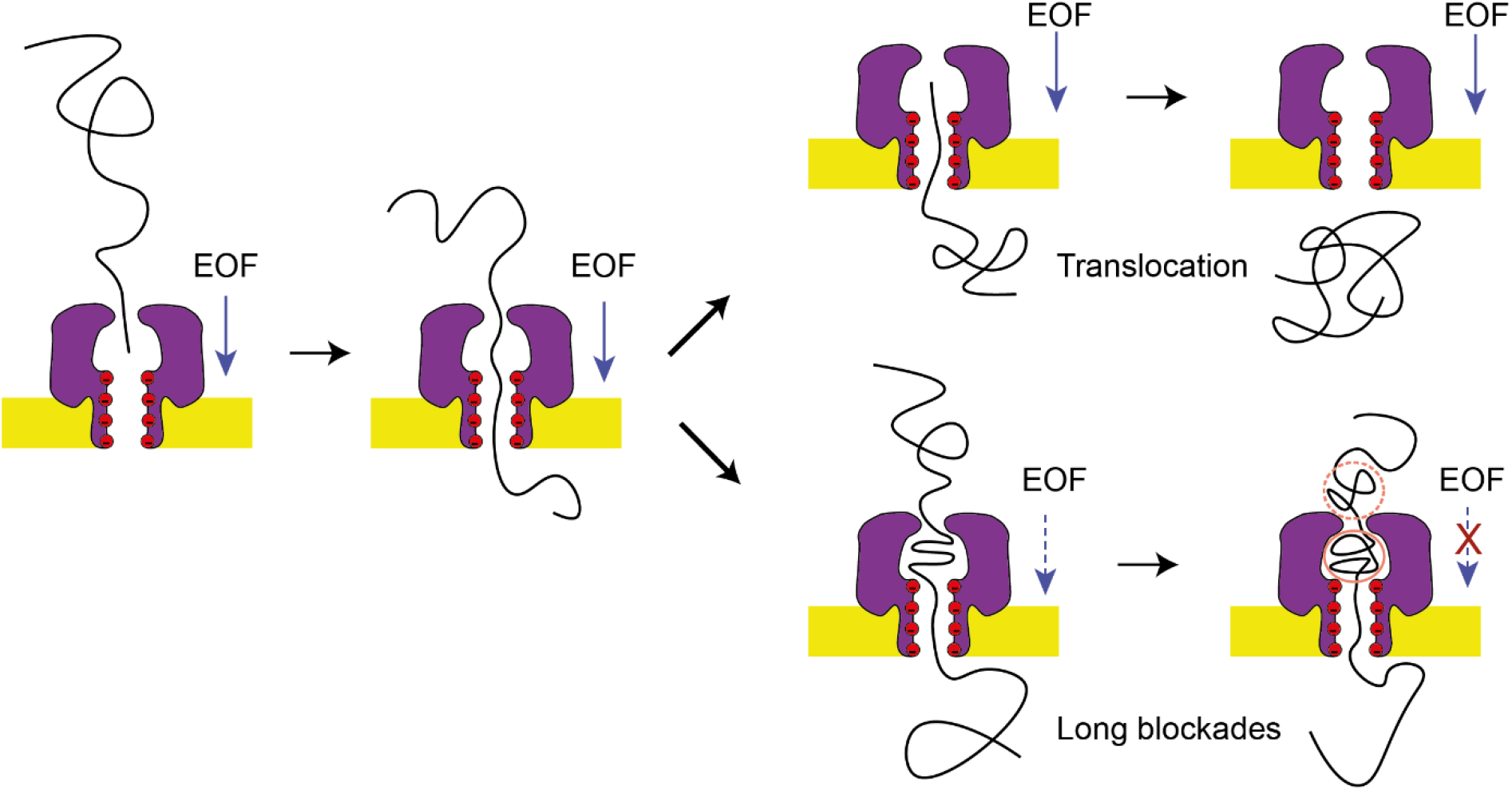
Potential mechanism for polypeptide translocation across CytK-4D at low potentials (< -120 mV) and high potentials (> -120 mV). In the higher regime, coils and blobs may start forming in the vestibule of the nanopore as the velocity of transport is increased.

### Role of the nanopore size, vestibule, and aromatic interactions in blob-like structure formation

The relevance of a narrow entry was probed by stepwise truncation of the first 8 N-terminal residues within CytK-4D, which define the entry of the nanopore (**Figure 4A, S1**). ΔN2, ΔN4 and ΔN6 truncations did not affect the translocation substantially (**Figures S13-S15**). On the other hand, ΔN8 truncation, which broadened the entry diameter to 2.7 nm, promoted the occurrence of long-lived multi-level events (**Figure S16**), compatible with the formation of blobs.

**Figure 4.**
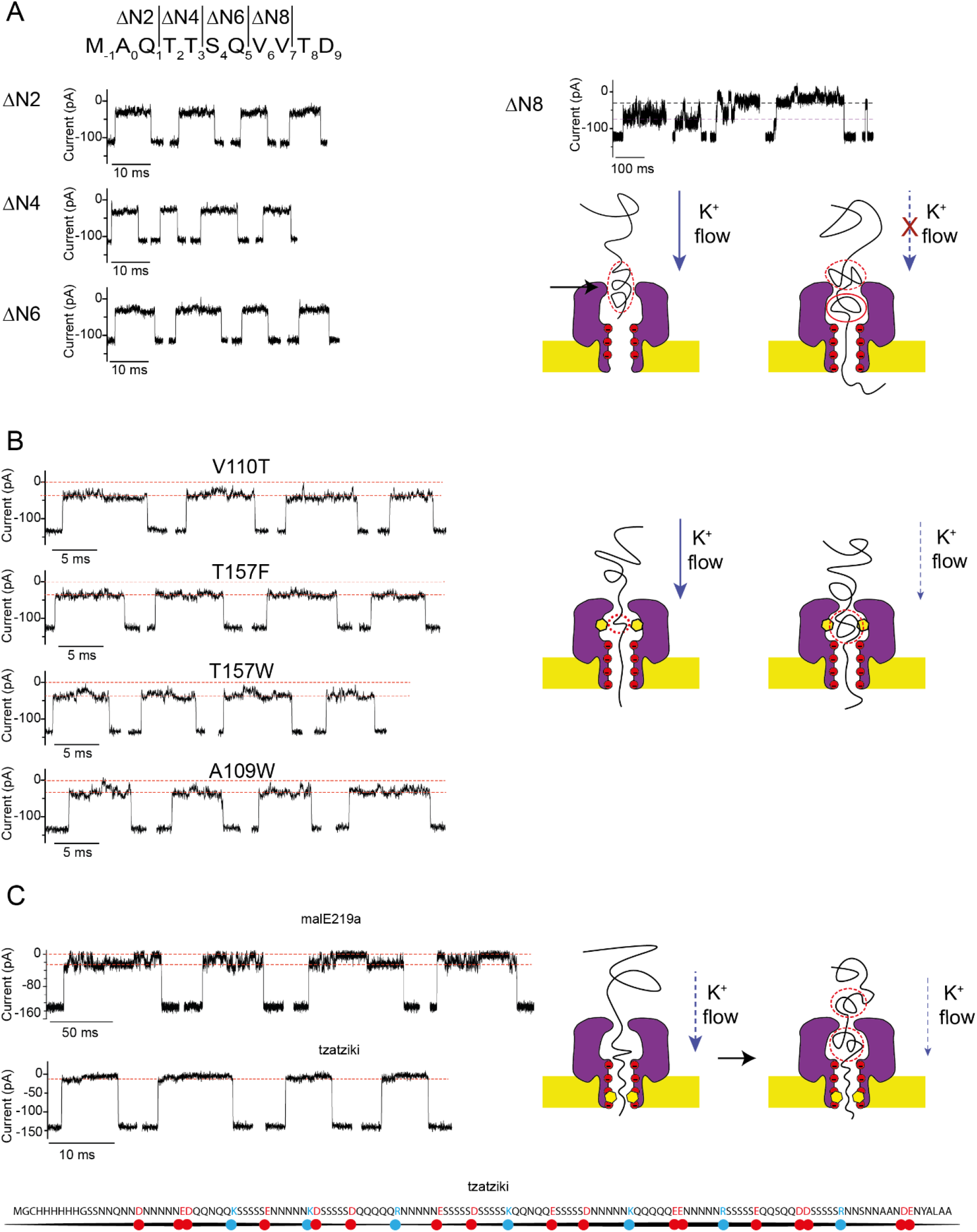
Mutagenesis in the CytK-4D nanopore addressing the formation of coil and blob-like structures within CytK-4D. A) Traces from the N-terminal truncation mutants and potential mechanism in the ΔN8 truncation. **B**) Traces from the vestibule mutants and potential mechanism. **C**) Traces from malE219a and tzatziki translocation through CytK-4D-S126F (-100 mV) and potential explanation for this behaviour. Data were collected in 1 M KCl, 2 M urea, 15 mM HEPES buffer pH 7.5. Traces were sampled at 50 kHz and filtered at 10 kHz.

If blobs / coils are formed inside the vestibule, amino acid substitutions are expected to affect the threading of substrates. The introduction of aromatic residues (A109W, T157F/W) or substitution of an aliphatic residue for a polar one (V110T) in the vestibule near the β-barrel entry (**Figure 4B, S17-S20)** resulted in longer events compared to CytK-4D (**Table S3**), showing that the polypeptide is likely to interact with the vestibule of CytK during translocation. This indicates that transient structures, such as coils or blobs, may form during translocation as the polypeptide samples the vestibule space of CytK prior to threading to the β-barrel region.

The role of polymer-pore interactions was further investigated by introducing an aromatic residue within the β-barrel of CytK-4D (CytK-4D-S126F, **Figure S21**). This substitution reduced the translocation velocity by ∼10 fold across all potentials (**Table S4**), and the I_ex%_ increased (e.g., to 89.2 ± 0.6 % compared to 71.3 ± 0.8 % for CytK-4D at -100 mV). Furthermore, current signatures similar to those obtained from MspA and lysenin nanopores were observed (**Figure 4C**), suggesting the formation of blob-like structures. A likely interpretation is that S126F leads to the coiling of the substrate within the vestibule by contributing to sudden changes in the local velocity once the polypeptide interacts with the introduced aromatic residue.

To confirm the role of aromatic interactions on blob-formation, we tested tzatziki, a model substrate devoid of aromatic residues (except for a single Tyr at the C-terminus) and rich in disorder-promoting residues such as Ser, Gly, Asn, and Gln, and charged residues (**Figure 4C**). Tzatziki was associated with a smooth translocation through CytK-4D (**Figure 4C**, I_ex%_ of ∼90 %, not affected by urea in solution). The presence of S126F did not alter the current signature of this substrate (**Figure S22)**, despite a slight increase in I_ex%_ (2-3 %, **Table S5**) and a ∼2-fold decreased velocity compared to CytK-4D. These observations indicate that polypeptide-nanopore interactions (*e*.*g*., through hydrophobic interactions) modulate the velocity and the formation of blob-like structures within the nanopore.

### Electrophoretic versus electroosmotic polymer translocation

The observation that linearised translocation can only be obtained under a narrow range of conditions is surprising. This is because the transport of ssDNA, which have a similar persistence length as polypeptides, is not associated to the formation of blob-like structures through a range of different conditions^1–3^. A likely interpretation lies in the fact that the DNA is uniformly charged, and its translocation is dominated by the electrophoretic force. Under electrophoretic transport, the electroosmotic component acts in the opposite direction of the polymer translocation and it contributes to stretching the polymer most likely preventing the formation of blob-like structures.

The translocation of two model unstructured polypeptides help to further shed light on the transport of polypeptides across nanopores ^7^. Highly positively charged S1, **Figure S23**), allowed us to test the electrophoretic-driven transport of a polypeptide. At translocation velocity of S1 across CytK-WT^7^ (EPF-driven transport) the long-lived events were absent, indicating that as for ssDNA an EOF opposing the direction of translocation prevents the formation of blob-like structures also with polypeptides. Weakly negatively charged tzatziki devoid of hydrophobic amino acids did not induce long-lived events, suggesting that under electroosmotic-driven transport blob-like structures in proteins are facilitated by aromatic/aliphatic interactions with the nanopore.

## Conclusions

The translocation of polypeptides across membranes is an important biological process that is not easily monitored by biophysical techniques. In biotechnology, the transport of unfolded polypeptides across nanopores under an applied potential is under investigation for identification and sequencing of proteins at the single-molecule level.
In this work, we found that during the unstructured translocation across nanopores of weakly charged polymers, blob-like structures likely form, and this process is augmented by interactions between the polypeptide and the nanopore. The formation of blob-like structures might be prevented by using nanopores having both a narrow entry and a narrow lumen (<∼1.5 nm), and by removing residues in the nanopore that might interact with the hydrophobic residues in the translocating polypeptide.

## Acknowledgments

The authors acknowledge the grants NWO-VICI (grant 192.068) and NIH (grant HG012554) and thank Gregor Anderluh for kindly providing the gene of the lysenin nanopore.

## References

1. Stoddart, D., Heron, A. J., Mikhailova, E., Maglia, G. & Bayley, H. Single-nucleotide discrimination in immobilized DNA oligonucleotides with a biological nanopore. Proceedings of the National Academy of Sciences 106, 7702–7707 (2009).

2. Butler, T. Z., Pavlenok, M., Derrington, I. M., Niederweis, M. & Gundlach, J. H. Single-molecule DNA detection with an engineered MspA protein nanopore. Proceedings of the National Academy of Sciences 105, 20647–20652 (2008).

3. Wloka, C., Mutter, N. L., Soskine, M. & Maglia, G. Alpha-Helical Fragaceatoxin C Nanopore Engineered for Double-Stranded and Single-Stranded Nucleic Acid Analysis. Angewandte Chemie International Edition 55, 12494–12498 (2016).

4. Yu, L. et al. Unidirectional single-file transport of full-length proteins through a nanopore. Nat Biotechnol (2023) doi:10.1038/s41587-022-01598-3.

5. Cressiot, B. et al. Dynamics and Energy Contributions for Transport of Unfolded Pertactin through a Protein Nanopore. ACS Nano 9, 9050–9061 (2015).

6. Pastoriza-Gallego, M. et al. Evidence of Unfolded Protein Translocation through a Protein Nanopore. ACS Nano 8, 11350–11360 (2014).

7. Sauciuc, A., Morozzo della Rocca, B., Tadema, M. J., Chinappi, M. & Maglia, G. Translocation of linearized full-length proteins through an engineered nanopore under opposing electrophoretic force. Nat Biotechnol (2023) doi:10.1038/s41587-023-01954-x.

8. Martin-Baniandres, P. et al. Enzyme-less nanopore detection of post-translational modifications within long polypeptides. Nat Nanotechnol (2023) doi:10.1038/s41565-023-01462-8.

9. Versloot, R. C. A., Straathof, S. A. P., Stouwie, G., Tadema, M. J. & Maglia, G. β-Barrel Nanopores with an Acidic–Aromatic Sensing Region Identify Proteinogenic Peptides at Low pH. ACS Nano 16, 7258–7268 (2022).

10. Manrao, E. A. et al. Reading DNA at single-nucleotide resolution with a mutant MspA nanopore and phi29 DNA polymerase. Nat Biotechnol 30, 349–353 (2012).

11. Brinkerhoff, H., Kang, A. S. W., Liu, J., Aksimentiev, A. & Dekker, C. Multiple rereads of single proteins at single–amino acid resolution using nanopores. Science (1979) 374, 1509–1513 (2021).

12. Chen, Z. et al. Controlled movement of ssDNA conjugated peptide through Mycobacterium smegmatis porin A (MspA) nanopore by a helicase motor for peptide sequencing application. Chem Sci 12, 15750–15756 (2021).

13. Yan, S. et al. Single Molecule Ratcheting Motion of Peptides in a Mycobacterium smegmatis Porin A (MspA) Nanopore. Nano Lett 21, 6703–6710 (2021).

14. Nova, I. C. et al. Detection of phosphorylation post-translational modifications along single peptides with nanopores. Nat Biotechnol (2023) doi:10.1038/s41587-023-01839-z.

15. Bogard, A. et al. The Ionic Selectivity of Lysenin Channels in Open and Sub-Conducting States. Membranes (Basel) 11, 897 (2021).

16. Ouldali, H. et al. Electrical recognition of the twenty proteinogenic amino acids using an aerolysin nanopore. Nat Biotechnol 38, 176–181 (2020).

17. Stirnemann, G., Giganti, D., Fernandez, J. M. & Berne, B. J. Elasticity, structure, and relaxation of extended proteins under force. Proc Natl Acad Sci U S A 110, (2013).

18. Gräter, F., Heider, P., Zangi, R. & Berne, B. J. Dissecting entropic coiling and poor solvent effects in protein collapse. J Am Chem Soc 130, (2008).

19. Cifra, P., Benková, Z. & Bleha, T. Persistence lengths and structure factors of wormlike polymers under confinement. Journal of Physical Chemistry B 112, (2008).

20. Vaitheeswaran, S. & Thirumalai, D. Interactions between amino acid side chains in cylindrical hydrophobic nanopores with applications to peptide stability. Proc Natl Acad Sci U S A 105, (2008).

21. Penkett, C. J. et al. Structural and Dynamical Characterization of a Biologically Active Unfolded Fibronectin-Binding Protein from Staphylococcus a ureus. Biochemistry 37, 17054–17067 (1998).

22. Bright, J. N., Woolf, T. B. & Hoh, J. H. Predicting properties of intrinsically unstructured proteins. Prog Biophys Mol Biol 76, 131–173 (2001).

23. Yang, G. et al. Solid-state synthesis and mechanical unfolding of polymers of T4 lysozyme. Proceedings of the National Academy of Sciences 97, 139–144 (2000).

24. Oesterhelt, F. et al. Unfolding Pathways of Individual Bacteriorhodopsins. Science (1979) 288, 143–146 (2000).

25. Carrion-Vazquez, M. et al. The mechanical stability of ubiquitin is linkage dependent. Nat Struct Mol Biol 10, 738–743 (2003).

26. Dietz, H. & Rief, M. Protein structure by mechanical triangulation. Proceedings of the National Academy of Sciences 103, 1244–1247 (2006).

27. Kohn, J. E. et al. Random-coil behavior and the dimensions of chemically unfolded proteins. Proceedings of the National Academy of Sciences 101, 12491–12496 (2004).

28. Daoud, M. & De Gennes, P. G. Statistics of macromolecular solutions trapped in small pores. Journal de Physique 38, 85–93 (1977).

29. Dai, L., Renner, C. B. & Doyle, P. S. The polymer physics of single DNA confined in nanochannels. Advances in Colloid and Interface Science vol. 232 Preprint at 10.1016/j.cis.2015.12.002 (2016).

30. Daoudi, S. & Brochard, F. Flows of Flexible Polymer Solutions in Pores. Macromolecules 11, 751–758 (1978).

31. Wong, C. T. A. & Muthukumar, M. Polymer capture by electro-osmotic flow of oppositely charged nanopores. J Chem Phys 126, (2007).

