## Supporting information for "Unravelled proteins form blobs during translocation across nanopores"

**Table of contents**

Materials and methods

Chemical reagents

Protein sequences nanopores

Cloning of nanopores

Expression and purification nanopores

Oligomerisation lysenin

Oligomerisation aerolysin

Substrates

Crystallisation

Electrophysiology measurements

Table S1. Data collection and refinement statistics (molecular replacement)

Table S2. Reversal potentials and cation selectivities of Aerolysin mutants

Table S3. Dwell times, $I_{ex\%}$ and relative velocity for the vestibule mutants

Table S4. Translocation malE219a through the CytK-4D-S126F mutant

Table S5. Translocation tzatziki though the CytK-4D-S126F mutant

Figure S1. Crystal structure of the CytK nanopore

Figure S2. Long-lived events occurring in translocation experiments

Figure S3. Long-lived events occurring in translocation experiments at low and high potential

Figure S4. Translocation malE219a through MspA WT nanopore

Figure S5. Translocation malE219a through lysenin WT nanopore

Figure S6. Addition of malE219a to Aerolysin mutant nanopores in cis

Figure S7. Addition of malE219a to Aerolysin 1D-3N in cis in higher salt

Figure S8. Addition of malE219a in trans

Figure S9. Translocation malE219a from trans to cis through the CytK-4D nanopore

Figure S10. Translocation malE219a from trans to cis through the Aer-1D-3N nanopore

Figure S11. Translocation malE219a from trans to cis through the Aer-2D-3N nanopore

Figure S12. Unstructured polymers in solution and in confinement

Figure S13. Translocation of malE219a through the CytK-4D-ΔN2 nanopore

Figure S14. Translocation of malE219a through the CytK-4D-ΔN4 nanopore

Figure S15. Translocation of malE219a through the CytK-4D-ΔN6 nanopore

Figure S16. Translocation of malE219a through the CytK-4D-ΔN8 nanopore

Figure S17. Translocation of malE219a through the CytK-4D-A109W nanopore

Figure S18. Translocation of malE219a through the CytK-4D-V110T nanopore

Figure S19. Translocation of malE219a through the CytK-4D-T157W nanopore

Figure S20. Translocation of malE219a through the CytK-4D-T157F nanopore

Figure S21. Translocation of malE219a through the CytK-4D-S126F nanopore

Figure S22. Translocation of tzatziki through the CytK-4D-S126F nanopore

Figure S23. Difference between EPF-driven and EOF-driven transport of S1

**Additional discussion**

**Materials and Methods**

*Chemicals and reagents*

The chemicals and suppliers used are listed as follows: Ampicillin sodium salt was purchased from Fisher Bio Reagents; chloramphenicol (≥98.0) from Sigma Life Science); urea (≥99.5%), guanidinium chloride (≥99.5%, biochemistry), Isopropyl β-D-thiogalactopyranoside (≥99.0%, dioxin-free, animal-free), LB medium, 2xYT medium, NaCl (≥99.5%), HEPES (PUFFERAN® CELLPURE® (≥99.5%), imidazole (≥99%), KCl (≥99.5%), Tris(2-carboxyethyl)phosphine hydrochloride (≥98.0%), Dodecyl-β-D-maltoside (≥99%) from Roth; and n-hexadecane (99% from Acros Organics; protease inhibitors (Pierce™ Protease inhibitor Mini tablets, EDTA-free); GeneJET gel extraction kit, GeneJET PCR purification kit, GeneJET Plasmid Miniprep kit were purchased from (Thermo Scientific); Ni-NTA agarose from Qiagen; Strep Tactin® Sepharose® and D-desthiobiotin from IBA Lifesciences; DPhPC from Avanti polar lipids, n-pentane from Sigma-Aldrich; Quick Start Bradford 1x Dye Reagent (Bio-Rad); DNA primers and gBlock™ from IDT. BL21(DE3) strain harbouring the pET-PfuX7 plasmid was kindly provided by prof. dr. Oscar Kuipers.

*Protein sequences of the nanopores*

- CytK WT:

MAQTTSQVVTDIGQNAKTHTSYNTFNNEQADNMTMSLKVTFIDDPSADKQIAVINTTGSFMKANPTLSDAPVDG
YPIPGASVTLRYPSQYDIAMNLQDNTSRFFHVAPTNAVEETTVTSSVSYQLGGSIKASVTPSGPSGESGATGQVTWS
DSVSYKQTSYKTNLIDQTNKHVKWNVFFNGYNNQNWGIYTRDSYHALYGNQLFMYSRTYPHETDARGNLVPMN
DLPTLTNSGFSPGMIAVVISEKDTEQSSIQVAYTKHADDYTLRPGFTFGTGNWVGNNIKDVDQKTFNKSFVLDWKN
KKLVEKKGSAHHHHHH

- MspA WT:

MGLDNELSLVDGQDRTLTVQQWDTFLNGVFPLDRNRLTREWFHSGRAKYIVAGPGADEFEGTLELGYQIGFPWSL
GVGINFSYTTPNILIDDGDITAPPFGLNSVITPNLFPGVSISADLGNGPGIQEVATFSVDVSGAEGGVAVSNAHGTVT
GAAGGVLLRPFARLIASTGDSVTTYGEPWNMNGSAGSAWSHPQFEK

- lysenin WT:

MSAKAAEGYEQIEVDVVAVWKEGYVYENRGSTSVDQKITITKGMKNVNSETRTVTATHSIGSTISTGDAFEIGSVEV
SYSHSHEESQVSMTETEVYESKVIEHTITIPPTSKFTRWQLNADVGGADIEYMYLIDEVTPIGGTQSIPQVITSRAKIIV
GRQIILGKTEIRIKHAERKEYMTVVSRKSWPAATLGHSKLFKFVLYEDWGGFRIKTLNTMYSGYEYAYSSDQGGIYFD
QGTDNPKQRWAINKSLPLRHGDVVTFMNKYFTRSGLCYDDGPATNVYCLDKREDKWILEVVGSGHHHHHH

- Aer-WT:

MAEPVYPDQLRLFSLGQGVCGDKYRPVNREEAQSVKSNIVGMMGQWQISGLANGWVIMGPGYNGEIKPGTAS
NTWCYPTNPVTGEIPTLSALDIPDGDEVDVQWRLVHDSANFIKPTSYLAHYLGYAWVGGNHSQYVGEDMDVTRD
GDGWVIRGNNDGGCDGYRCGDKTAIKVSNFAYNLDPDSFKHGDVTQSDRQLVKTVVGWAVNDSDTPQSGYDV
TLRYDTATNWSKTNTYGLSEKVTTKNKFKWPLVGETELSIEIAANQSWASQNGGSTTTSLSQSVRPTVPARSKIPVKI
ELYKADISYPYEFKADVSYDLTLSGFLRWGGNAWYTHPDNRPNWNHTFVIGPYKDKASSIRYQWDKRYIPGEVKW

WDWNWTIQQNGLSTMQNNLARVLRPVRAGITGDFSAESQFAGNIEIGAPVPLAADSKVRRARSVDGAGQGLRLE
IPLDAQELSGLGFNNVSLSVTPAANQGSSHHHHHH

*Cloning of the nanopores*

Plasmids containing the mutant CytK nanopores were constructed by means of USER cloning[1,2]. PfuX7 DNA polymerase was prepared as previously described[3] and used to generate the fragments used in the USER reaction. Homology regions of 8-13 bps were defined in the vicinity of the ATG codon, the mutation site and downstream the stop codon, to mediate the joining of the empty vector with the gene fragments. The full insert was divided into upstream and downstream fragments at the mutation site and amplified with uracil-containing primers. The empty pT7-SC1 backbone (AmpR) was separately linearised using uracil-containing primers. The PCRs were performed as in ref[3], with the exception of the extension time, which was decreased from 1 min/kb to 30 s/kb. The PCR products were purified either by gel extraction (if by-products were present), or by cleaned up from the PCR mix. The upstream (F1) and downstream (F2) fragments were added to the empty vector (V) in a molar ratio F1:F2:V of 3:3:1 in the USER reaction, which was carried out as in ref[2]: 25 min at 37 °C, followed by 10 min incubation at 60 °C. Lastly, the mixture was cooled down to room temperature (22 or 20 °C) for 15 min and subsequently stored on ice/ at 4 °C until the transformation step. The annealed fragments resulting in circularised plasmids were transformed into chemically (RbCl protocol) competent *E. coli* cells using the heat shock procedure (42 °C for 70 s followed by incubation on ice for 10 min) and the cells were selected on a LB-agar plate supplemented with 100 µg/mL ampicillin and 1% glucose. Plasmids from individual colonies were isolated and the introduction of the mutations was confirmed by Sanger sequencing (Macrogen/ Eurofins).

pT7-SC1 bearing MspA-WT was generated from pT7-SC1 by means of fusion PCR followed by USER cloning. First, three gene fragments were generated, with the overlap regions for fusion PCR being in the vicinity of the 90-93 residues and the 134-139 residues. In the second PCR round, the three fragments were joined together in the presence of U-containing primers. This insert and the empty pT7-SC1 vector were used for USER cloning.

pT7-SC1 bearing pro-aerolysin K238D, previously prepared in our laboratory[4], was used to generate the 1D-3N and 2D-3N aerolysin mutants. The mutations, namely R220N, R282N, K242N and S272D were introduced as described for the CytK mutants.

pT7-SC1 bearing lysenin-WT was previously prepared in our laboratory from the plasmid kindly provided by Gregor Anderluh.

*Expression and purification of the nanopores*

The plasmids encoding for the nanopores were electroporated into BL21(DE3) electrocompetent cells using a Bio Rad Micro Pulser (bacterial setting). The cells were selected on plates supplemented with 100 µg/mL ampicillin and 1% glucose and grown at 37 °C overnight. Next, several transformants were resuspended in 200 mL (proaerolysin, lysenin, MspA) or 100 mL (CytK) LB medium containing 100 µg/mL ampicillin, such that the starting optical density at 600 nm ($OD_{600}$) was 0.05-0.1. Cells were grown at 37 °C, 180 RPM until an $OD_{600}$ of 0.6-0.8. At this point, protein expression was induced with 0.5 mM IPTG after the cultures were chilled on ice for 5-10 min. The cells were harvested (7500 rpm, 5 min) following an incubation of 19-21 h at 25 °C, 180 RPM. The resulting cell pellets were incubated at least 30 min at -80 / -70 °C, prior to purification. Pellets from 100 mL cultures were resuspended in 20-25 mL ice-cold lysis buffer, while pellets from 200 mL cultures were resuspended in 30-40 mL lysis buffer. Subsequent steps were performed at 4-6 °C, unless stated otherwise. The cell suspension was sonicated with a Branson sonifier 450 at 25% duty cycle, 2.5 output control for 2-3 min in the case of the CytK mutants, or with a Branson sonifier 550 using pulses at 15 % amplitude, 5 s on, 7 s off, total 5 min on time in the case of MspA, lysenin and aerolysin mutants. Following sonication, the cellular debris was removed (8000 RPM, 20 min). The resulting supernatant was incubated with 200 µL $Ni^{2+}$-NTA slurry (50 % suspension) in the case of CytK, Aer and lysenin, or StrepTactin slurry in the case of MspA, pre-equilibrated and prewashed with 1 mL lysis buffer, for 20-40 min with shaking. The beads were briefly pelleted (1500 RPM, 1 min) and transferred to the column (2 mL bed volume bio-spin chromatography, BioRad) while allowing the flow through pass, at RT. The column was washed in steps with a total of 10 mL wash buffer. The protein was eluted with 150 µL elution buffer in three elution fractions. The presence of the SDS-stable CytK mutant oligomers and MspA oligomers was confirmed by SDS-PAGE, omitting the heating step in the sample preparation.

*Purification buffers*

| Nanopore | Lysis buffer | Wash buffer | Elution buffer |
|---|---|---|---|
| CytK | 50 mM HEPES, 150 mM NaCl, 10 mM imidazole, pH 7.4 + 0.02% DDM | 50 mM HEPES, 150 mM NaCl, 30 mM imidazole, pH 7.4 + 0.02% DDM | 50 mM HEPES, 150 mM NaCl, 250 mM imidazole, pH 7.4 + 0.02% DDM |
| MspA | 50 mM HEPES, 150 mM NaCl, pH 7.4 + 0.02% DDM | 50 mM HEPES, 150 mM NaCl, pH 7.4 + 0.02% DDM | 50 mM HEPES, 150 mM NaCl, pH 7.4 + 0.02% DDM + 3 mM desthiobiotin |
| Lysenin | 50 mM HEPES, 150 mM NaCl, 10 mM imidazole, pH 7.4 | 50 mM HEPES, 150 mM NaCl, 30 mM imidazole, pH 7.4 | 50 mM HEPES, 150 mM NaCl, 300 mM imidazole, pH 7.4 |
| Aerolysin | 50 mM HEPES, 500 mM NaCl, 10 mM imidazole, pH 7.4 | 50 mM HEPES, 500 mM NaCl, 30 mM imidazole, pH 7.4 | 50 mM HEPES, 500 mM NaCl, 250 mM imidazole, pH 7.4 |

<u>*Oligomerisation lysenin*</u>

The concentration of the elution fractions was determined by Bradford assay. Sphingomyelin/DPhPC liposomes (50/50) were added to 0.25 mg/mL lysenin sample in a 10:1 w/w ratio, then incubated at 37 $^{o}$C for 30 min. The lysenin oligomers were extracted from the liposomes by adding 6 % LDAO (in MQ) in a 1:10 v/v ratio to the protein-liposome solution. Next, the mixture was further diluted ~25-fold in 150 mM NaCl, 15 mM Tris + 0.02 % DDM (total final volume 6 mL). 50 µL pre-equilibrated $Ni^{2+}$-NTA beads were added and the solution was incubated at room temperature for 10 min with shaking. The beads were transferred to the column, washed with a total of 10 mL 150 mM NaCl, 15 mM Tris + 0.02 % DDM to remove the LDAO and finally eluted with 100 µL EBO buffer (15 mM TrisHCl pH 7.5, 150 mM NaCl, 200 mM $Na_2$EDTA pH 8.0, 0.02 % DDM). The oligomers were stored at 4 $^{o}$C for several months.

<u>*Oligomerisation aerolysin*</u>

The pro-aerolysin elution fractions were combined and subsequently desalted by using an amicon filter 10 K MWCO and 50 mM HEPES, 150 mM NaCl, pH 7.4 buffer. This sample was used for oligomerisation. To promote its oligomerisation, the proaerolysin protein (~0.5 mg/mL) was cleaved by adding 0.01

mg/mL trypsin in a proaerolysin: trypsin volume ration 10:1. Additionally, 0.02 % DDM was added during cleavage, in order to improve the stability of the oligomers. This mix was incubated at room temperature for 20 min, then stored at 4 °C for 2-3 weeks.

*Substrates*

The protein substrates were prepared as previously described[5].

*Crystallisation of the CytK nanopore*

For crystallisation and structure determination, we selected the CytK 2E-2D-T147D mutant due to its improved solubility compared to any other mutant. pT7-SC1 harbouring this CytK mutant was electroporated into electrocompetent *E. coli* BL21(DE3) and transformants were selected on LB-agar plates supplemented with 100 µg/mL ampicillin and 1% glucose (37 °C overnight). 800 mL culture was grown and the mutants was purified as described in the previous section.

Following size exclusion chromatography, the protein was kept in a solution of 50 mM HEPES, pH 7.4, 150 mM NaCl and 0.02 % DDM which was subsequently concentrated to 8 mg/mL using a 30,000 MWCO concentrator. CytK was mixed with the crystallisation buffer in a ratio of 2:1 and all crystals were obtained via sitting drop vapour diffusion using Molecular Dimensions screening kits (Calibre Scientific). Crystal hits were observed in numerous conditions using the MemGold I and Morpheus II screening kits, the crystals which appeared the most optimal were further optimised with the custom made screens made with the DragonFly (SPT Labtech). The optimised conditions produced yellow, rectangular prismatic crystals of approximately 500 microns in length along the longest axis in the following condition: 0.005 M Manganese(II) chloride tetrahydrate, 0.005 M Cobalt(II) chloride hexahydrate, 0.005 M Nickel(II) chloride hexahydrate, 0.005 M Zinc acetate dihydrate, 0.1 M MOPSO, Bis-Tris, pH 6.5, 35 %(w/v) glycerol, 30 %(w/v) PEG 4000. Crystals were briefly soaked in 50 %(w/v) PEG 4000 for cryo-protection and subsequently flash-frozen in liquid nitrogen in preparation for diffraction experiments at a synchrotron. Data were collected at beam line ID23-1 (ESRF, Grenoble).

The most optimal crystal of CytK diffracted at 4.3 Å resolution (Table S1). Data were processed with XDS[6] and structures were solved by Molecular Replacement with Phaser[7] using a previously published homologous model of alpha hemolysin (PDB ID: 7AHL). Manual rebuilding was performed with COOT[8] and refinement with PHENIX[9]. In all chains, minor truncation of small amino acid strings was necessary as there was no assignable electron density nearby. Data collection and refinement statistics are given in Table S1. The refined model was deposited into the PDB repository with accession code 8RJ8. Images were prepared using UCSF ChimeraX[10].

*Electrophysiology measurements*

Recordings in planar lipid bilayers were carried out using a chamber comprising of two compartments, separated by a 25 µm thick Teflon membrane containing an aperture of approximately 100 µm as described earlier[11]. A droplet (half the quantity contained in a 10 µL glass capillary) consisting of n-hexadecane dissolved in n-pentane (6.25%) was applied on the Teflon membrane. When the n-pentane evaporated, 500 µL buffer, followed by two droplets of DPhPC lipids in n-pentane (5 mg/mL) were added in each compartment. Ag/AgCl electrodes were connected to the two compartments via agarose bridges (2.5% agarose, 3M KCl solution), grounding the *cis* compartment. Measurements were performed using an Axon™ Digidata® 1550B digitizer and an Axopatch 200B amplifier (Molecular Devices) and recorded with the Clampex 11.1 software.

*Ion selectivity*

Buffers:  Buffer A - 2 M KCl, 15 mM HEPES, pH 7.5, Buffer B - 0.5 M KCl, 15 mM HEPES, pH 7.5, Buffer C - 0 M KCl, 15 mM HEPES, pH 7.5,

The ion selectivity of the aerolysin mutants and MspA WT was determined using the buffers described above. Firstly, both the *cis* and *trans* compartments were filled with 500 µL buffer A and a single nanopore was isolated. Next, the pipet-offset was adjusted to 0 pA at 0 mV bias. The I/V curve was determined between -30 and +40 mV, in 5 mV steps, using a 10 kHz sampling rate coupled with a 2 kHz Bessel filter. Next, the KCl concentration in the *trans* compartment was decreased to approximately 0.5 M by perfusing with buffer C (5 x 100 µL). Subsequent perfusion with buffer B (7 x 100 µL) ensured that the final concentration of KCl in the *trans* compartment was correctly fine-tuned to 0.5 M. A similar protocol was used to record the I/V curve of the nanopore in these conditions. This second I/V curve was used to determine the reversal potential from the linear function fitting the data points between -20 and +20 mV. Finally, the ion selectivity, expressed as the fraction $p_{K+}/p_{Cl-}$ was calculated using the formula below and each ion selectivity was established from triplicate experiments.

$$\frac{p_{K+}}{p_{Cl-}} = \frac{[a_{Cl^-}]_{trans} - [a_{Cl^-}]_{cis} \times e^{V_r F/RT}}{[a_{K^+}]_{trans} \times e^{V_r F/RT} - [a_{K^+}]_{cis}}$$

where [a] is the activity of the $K^+$ or $Cl^-$ in the *cis* or trans compartment, $V_r$ is the reversal potential, which is obtained from the experiments, F corresponds to the Faraday constant (96 485 C/mol), R the gas constant (8.3145 J $mol^{-1}$ $K^{-1}$) and T the temperature (298 K).

*Translocation experiments*

Single pores were isolated and the pore orientation was determined based on the I/V curve (1 M KCl, 15 mM HEPES, pH 7.5). 2 M urea was introduced into the system by exchanging 125 µL buffer with 125 µL 1 M KCl, 8 M urea, 15 mM HEPES, pH 7.5 in both compartments. Next, 0.3 µM malE219a was added in the *cis* compartment and translocation was induced by applying negative potential. Alternatively, the protein was added in the *trans* compartment and events were induced by applying positive potential. Each nanopore (substrate in either cis or trans) was tested in triplicate.

*Data analysis translocation experiments*

The generated files were analysed using the Clampfit 11.1 software. Firstly, the open pore current (level 0, L0) and the corresponding noise, σ, were obtained from a full point histogram. Secondly, L1, corresponding to the detection limit, was set at 10σ, (note that Clampfit sets its half-way detection algorithm at 5σ). Events were recorded using voltage protocols in which the potential was ramped through positive and negative applied potentials. In the case of the *cis*-to-trans translocation, events were detected at negative potentials from approximately 2 s of recording times at negative bias. The resulting L1 data points were used to construct the log(dwell time) vs amplitude scatter plot, from which the amplitude boundaries of the event cluster were defined. Events faster than 100 µs were ignored. Next, using the amplitude boundaries, the logarithmic histogram of the dwell time and the conventional histogram of either the amplitude or the $I_{ex\%}$ were constructed. In both cases, the bin value was set such that the distribution within the histogram would resemble a Gaussian distribution as much as possible. The values for the log(dwell time) and either the amplitude or the $I_{ex}$ were established by fitting a Gaussian function to the histogram, whose μ is either log(dwell time) or the amplitude/$I_{ex\%}$. The dwell time and $I_{ex\%}$ are reported from triplicate measurements, with the values representing the average obtained from three individual repeats and the corresponding standard deviation (SD).

List of Tables

*Table S1. Data collection and refinement statistics (molecular replacement)*

| | ***CytK*** |
|---|---|
| **PDB-ID** | **XXXX** |
| Wavelength | 0.8856 |
| Resolution range | 88.02 - 4.32 (4.474 - 4.32) |
| Space group | P 1 21 1 |
| *a*, *b*, *c* (Å) | 103.229 186.222 104.532 |
| α, β, γ (°) | 90 115.85 90 |
| Unique reflections | 23236 (1713) |
| Completeness (%) | 95.69 (72.22) |
| Mean I/sigma(I) | 9.2 (4.61) |
| Wilson B-factor | 83.99 |
| R-meas | 0.16 |
| CC1/2 | 95.2 (25.0) |
| Reflections used in refinement | 22871 (1713) |
| Reflections used for R-free | 1262 (84) |
| R-work | 0.3254 (0.2715) |
| R-free | 0.3894 (0.3153) |
| Number of non-hydrogen atoms | 15699 |
| macromolecules | 15699 |
| ligands | 0 |
| solvent | 0 |
| Protein residues | 2012 |
| RMS(bonds) | 0.007 |
| RMS(angles) | 1.20 |
| Ramachandran favored (%) | 82.79 |
| Ramachandran allowed (%) | 13.79 |
| Ramachandran outliers (%) | 3.42 |
| Rotamer outliers (%) | 0.61 |
| Clashscore | 27.50 |
| Average B-factor | 89.23 |
| macromolecules | 89.23 |

*Values in parentheses are for the highest resolution shell.

*Table S2. Reversal potentials and cation selectivities of Aerolysin mutants*

| **Mutant** | **Vr** | **pK/pCl** |
|---|---|---|
| Aer K238D R220N | 3.22 ± 0.04 | 1.26 ± 0.004 |
| Aer K238D R282N | 2.76 ± 0.34 | 1.22 ± 0.04 |
| Aer K238D K242N | 9.81 ± 0.27 | 2.06 ± 0.05 |
| Aer K238D K242D | 9.99 ± 0.10 | 2.08 ± 0.02 |
| Aer K238D R220N R282N K242N (Aer 1D-3N) | 17.39 ± 0.16 | 3.95 ± 0.08 |
| Aer K238D R220N R282N K242N S272D (Aer 2D-3N) | 19.04 ± 0.31 | 4.68 ± 0.19 |

*Table S3. Dwell times, $I_{ex}$ and relative velocity (ratio between point mutation mutant and the original 4D nanopore, expressed as percentage) and the corresponding standard deviations for the vestibule mutants.*

| | | | | | | |
|---|---|---|---|---|---|---|
| 4D | | | | | | |
| V (mV) | time (ms) | stdev time | Iex (%) | stdev Iex | relative velocity (%) | stdev relative velocity |
| -60 | 24.44 | 6.73 | 72.29 | 1.32 | 100.00 | 20.72 |
| -80 | 10.92 | 0.65 | 71.86 | 0.76 | 100.00 | 6.19 |
| -100 | 5.51 | 0.45 | 71.27 | 0.79 | 100.00 | 8.14 |
| -120 | 3.28 | 0.45 | 70.06 | 0.57 | 100.00 | 12.75 |
| A109W | | | | | | |
| V (mV) | time (ms) | stdev time | Iex (%) | stdev Iex | relative velocity (%) | stdev relative velocity |
| -60 | 39.87 | 8.55 | 75.02 | 0.61 | 60.60 | 14.66 |
| -80 | 11.40 | 0.22 | 73.72 | 0.86 | 95.49 | 1.79 |
| -100 | 6.98 | 0.77 | 72.79 | 0.14 | 79.21 | 9.14 |
| -120 | 5.42 | 0.30 | 71.73 | 0.71 | 59.93 | 3.24 |
| -140 | 3.54 | 0.27 | 71.60 | 0.30 | | |
| T157F | | | | | | |
| Voltage | time (ms) | stdev time | Iex (%) | stdev Iex | relative velocity (%) | stdev relative velocity |
| 60 | 27.34 | 1.61 | 73.62 | 0.58 | 85.51 | 4.88 |
| -80 | 10.82 | 0.52 | 73.23 | 0.49 | 100.75 | 4.97 |
| -100 | 7.03 | 0.41 | 71.88 | 0.49 | 78.11 | 4.36 |
| -120 | 4.53 | 0.37 | 70.27 | 0.66 | 71.98 | 6.08 |
| T157W | | | | | | |
| V (mV) | time (ms) | stdev time | Iex (%) | stdev Iex | relative velocity (%) | stdev relative velocity |
| -60 | 28.47 | 3.40 | 73.95 | 1.51 | 82.85 | 10.14 |
| -80 | 11.11 | 1.43 | 73.32 | 1.27 | 99.27 | 13.59 |
| -100 | 6.99 | 0.23 | 71.87 | 0.60 | 78.49 | 2.54 |
| -120 | 4.46 | 0.43 | 70.62 | 0.62 | 73.12 | 6.90 |
| -140 | 3.27 | 0.56 | 68.98 | 0.58 | | |
| V110T | | | | | | |
| V (mV) | time (ms) | stdev time | Iex (%) | stdev Iex | relative velocity (%) | stdev relative velocity |
| -60 | 29.87 | 5.23 | 73.36 | 0.78 | 79.92 | 14.01 |
| -80 | 10.85 | 2.88 | 72.81 | 0.17 | 89.27 | 6.69 |
| -100 | 6.77 | 0.07 | 72.17 | 0.50 | 81.02 | 0.86 |
| -120 | 4.40 | 0.90 | 69.42 | 3.11 | 75.85 | 16.21 |

Table S4 Translocation malE219a through the CytK-4D-S126F mutant

| V (mV) | time (ms) | Iex (%) | velocity (nm/ms) | stdev time | stdev Iex | stdev velocity | relative velocity (%) | stdev relative velocity |
|---|---|---|---|---|---|---|---|---|
| -60 | 210.73 | 88.27 | 0.67 | 14.77 | 0.17 | 0.05 | 11.11 | 0.79 |
| -80 | 83.70 | 88.67 | 1.68 | 6.40 | 0.26 | 0.13 | 13.06 | 1.00 |
| -100 | 52.32 | 89.23 | 2.72 | 2.60 | 0.63 | 0.11 | 10.66 | 0.44 |
| -120 | 40.17 | 90.53 | 3.51 | 4.63 | 0.00 | 0.40 | 8.12 | 0.94 |
| -140 | 29.01 | 90.71 | 4.83 | 0.24 | 0.51 | 0.04 | | |

*Table S5. Translocation tzatziki though the CytK-4D-S126F mutant*

| V (mV) | time (ms) | stdev t | ln[t(ms)] | stdev ln t | Iex (%) | stdev Iex | relative velocity (%) | stdev relative velocity |
|---|---|---|---|---|---|---|---|---|
| -100 | 18.17 | 12.37 | 2.76 | 0.63 | 96.45 | 0.56 | 34.39 | 17.02 |
| -120 | 4.03 | 0.72 | 1.38 | 0.17 | 95.45 | 0.63 | 43.78 | 7.11 |
| -140 | 1.64 | 0.25 | 0.49 | 0.16 | 93.86 | 0.16 | 64.81 | 10.84 |
| -160 | 1.03 | 0.09 | 0.02 | 0.08 | 92.61 | 0.12 | 64.15 | 5.32 |
| -180 | 0.77 | 0.07 | -0.26 | 0.09 | 91.68 | 0.04 | 53.79 | 4.67 |

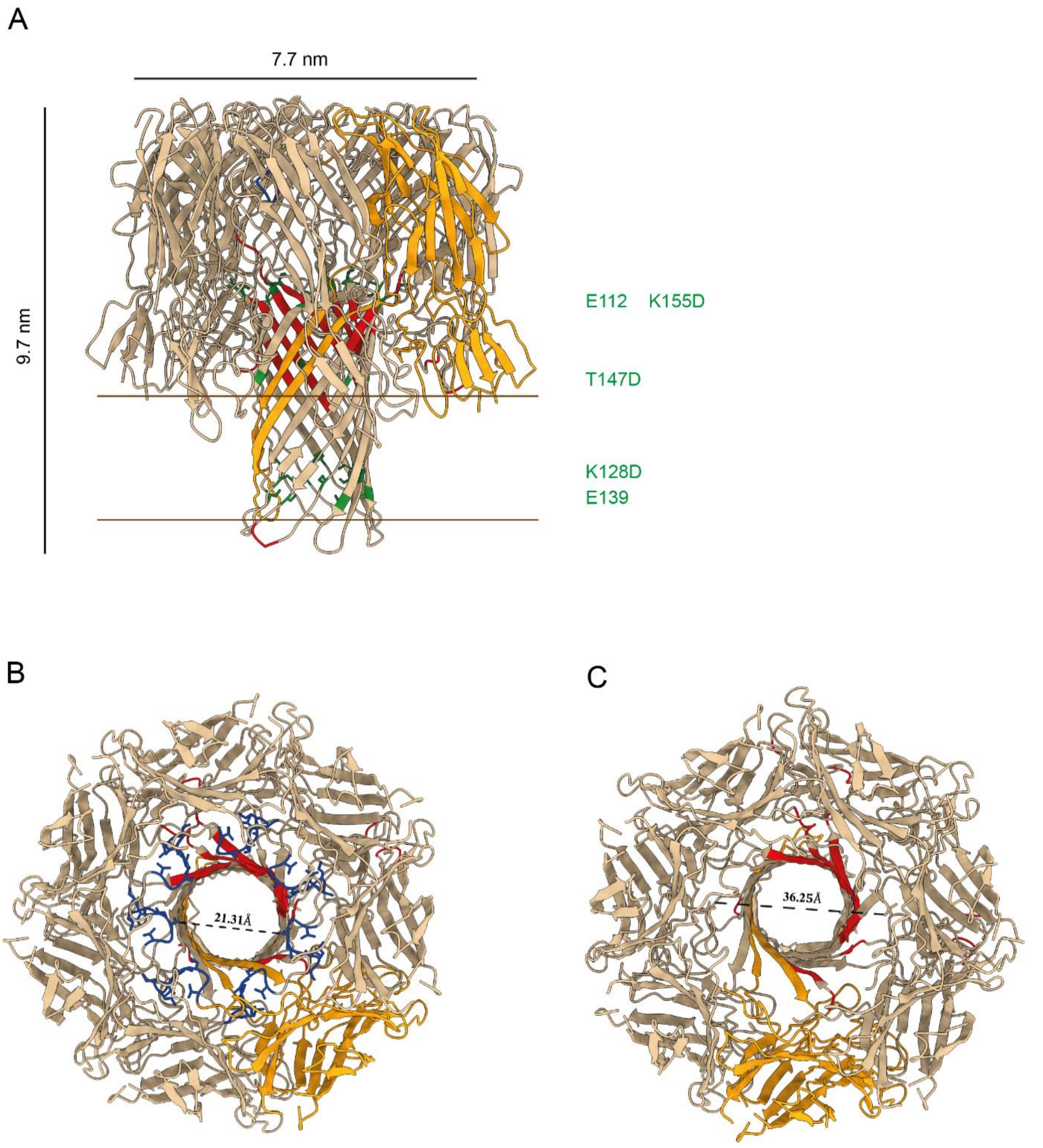

***Figure S1. Crystal structure of the CytK nanopore**, PDB 8RJ8. **A**) Structure of the CytK nanopore. Residues generated by NCS due to low electron density are depicted in red; protomer in bright orange and the negatively charged residues within the β-barrel in green. **B**) Top view of the nanopore, with $A_0$-$V_7$ highlighted in blue. **C**) Top view of the nanopore with $A_0$-$V_7$ removed.*

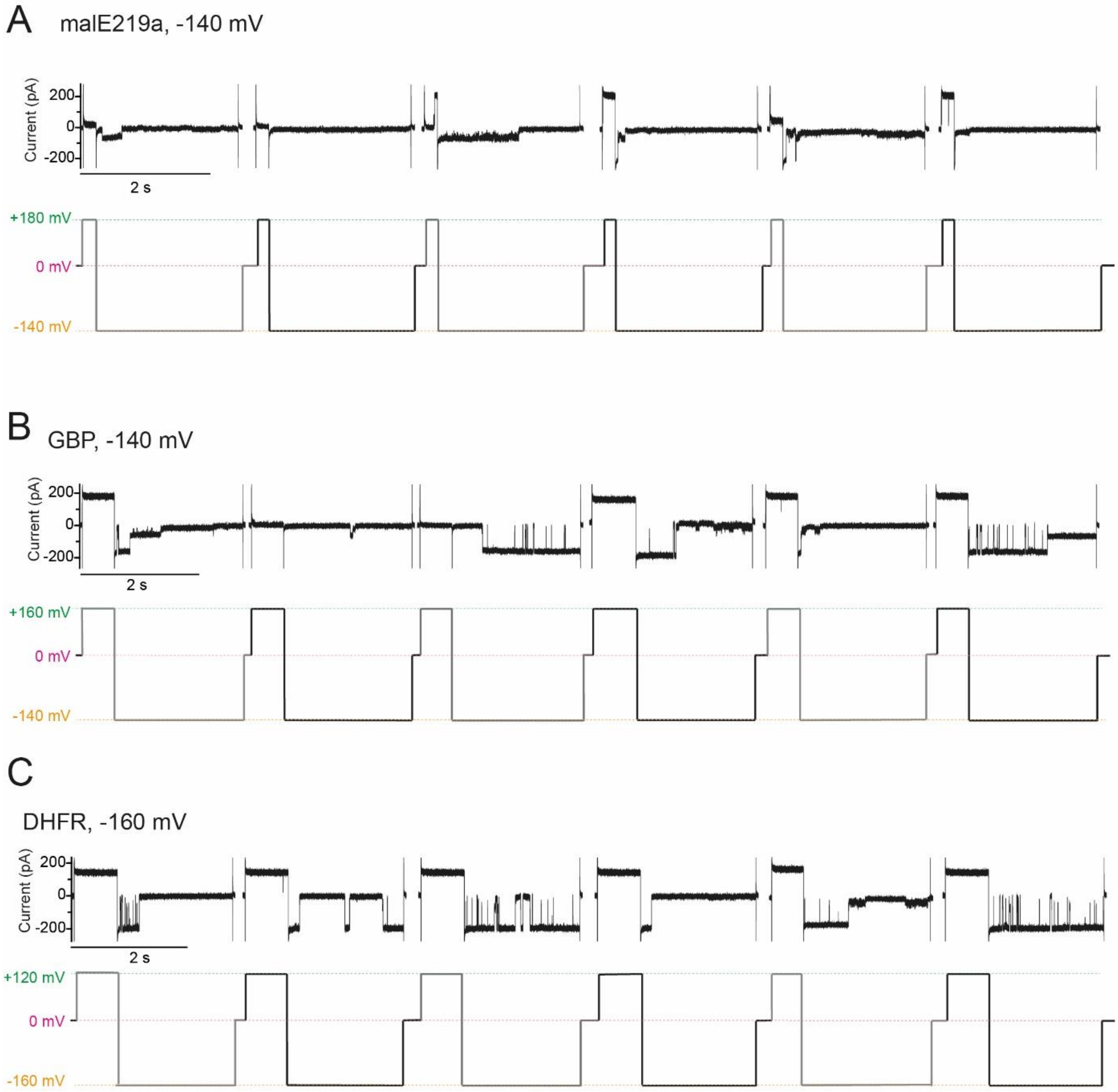

***Figure S2*. *Long-lived events occurring in translocation experiments carried out at high potentials in the case of the native substrates.*** *Each panel consists of traces (top) and the corresponding applied potentials (bottom). Substrates were added in cis and all traces were obtained in 1 M KCl, various urea concentrations (see later), 15 mM HEPES, pH 7.5, 50 kHz sampling and 10 kHz Bessel filter.* ***A****) malE219a at -140 mV (2 M urea).* ***B****) GBP H152A at -140 mV (2.4 M urea).* ***C****) DHFR W30G W133L at -160 mV (2.6 M urea).*

`

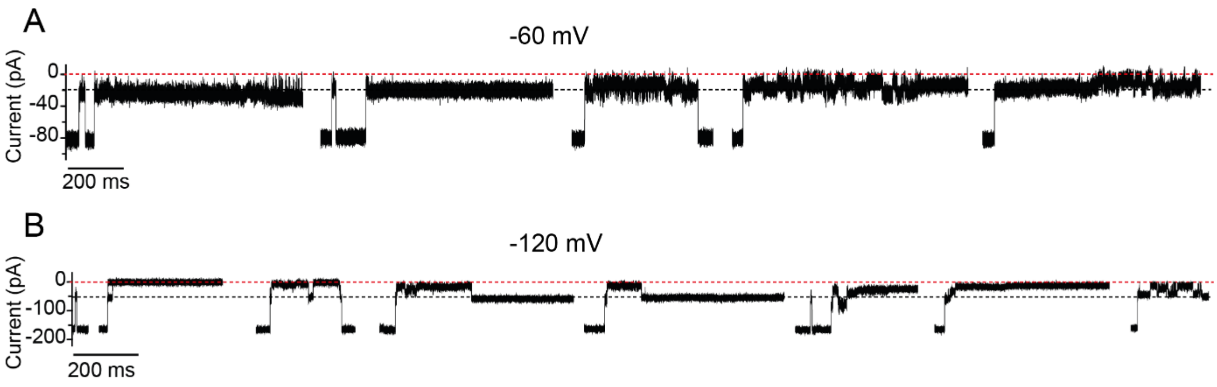

***Figure S3**. **Long-lived events occurring in translocation experiments. A**) Prolonged events at -60 mV and at -120 mV (**B**). The nanopore was CytK-4D nanopore and the polypeptide malE219a added in cis in 1 M KCl, 2 M urea, 15 mM HEPES, pH 7.5. Traces were collected at 50 kHz sampling rate and 10 kHz Bessel filter.*

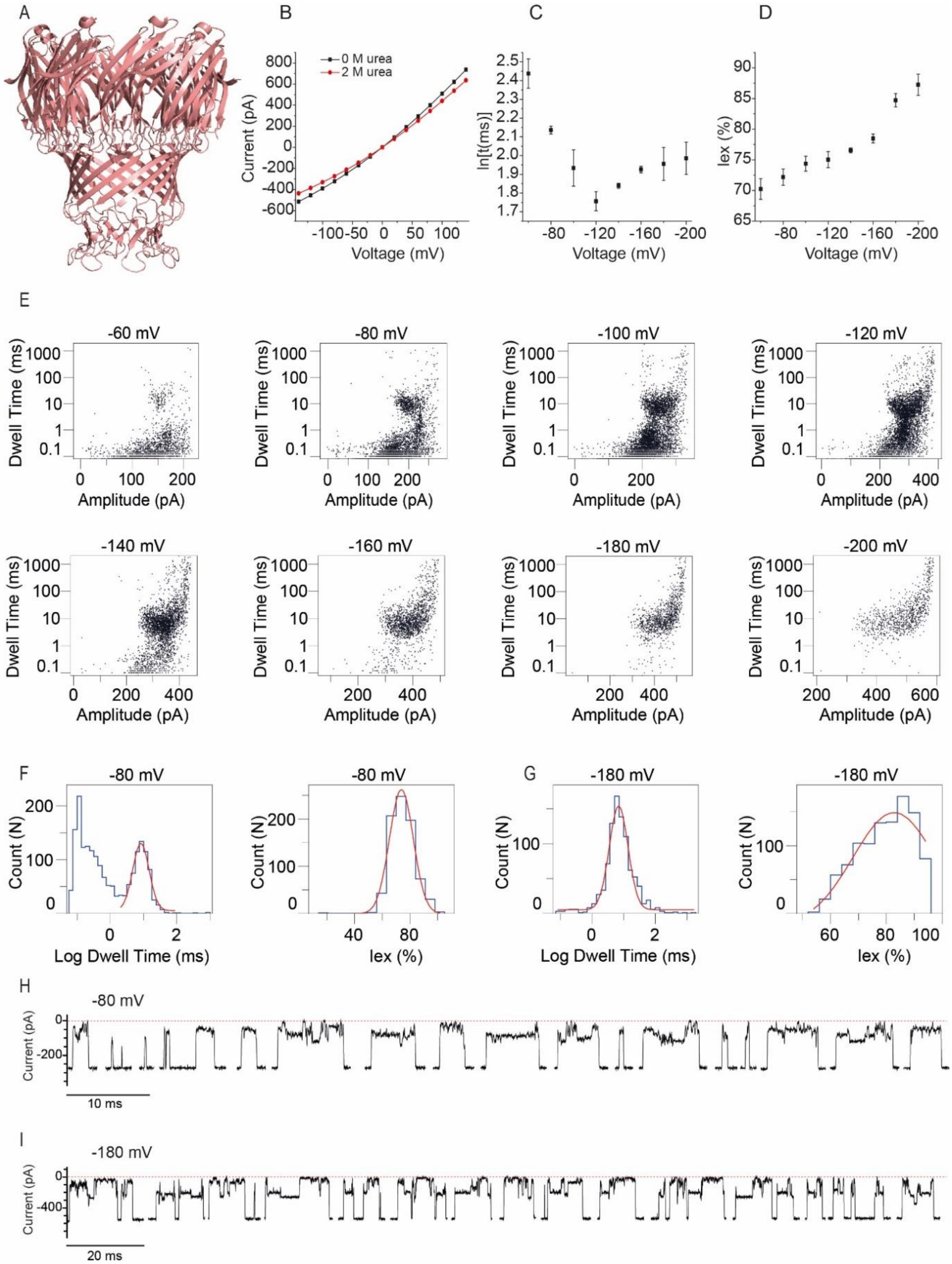

***Figure S4. Translocation of malE219a through MspA-WT nanopore. A)*** *Cartoon representation of the MspA nanopore.* ***B)*** *I-V curves in 1 M KCl at pH 7.5 of the MspA nanopore in 0 M and 2 M urea.* ***C)*** *Dwell time dependence on the applied potential for type I blockades.* ***D)*** *$I_{ex\%}$ dependence on the applied potential. All data points represent the average of three independent experiments and the error bars corespond to standard deviation (SD).* ***E)*** *Scatter plots from -60 to -200 mV.* ***F-G)*** *Examples of histograms for the log(dwell time) and $I_{ex\%}$ at -80 mV (F) and -180 mV (G).* ***H-I)*** *Typical events at -80 mV (H) and -180 mV, respectively (I). Recordings were performed in 1 M KCl, 2 M urea, 15 mM HEPES, pH 7.5, 50 kHz sampling, 10 kHz Bessel filter. Each measurement was performed in a least a triplicate.*

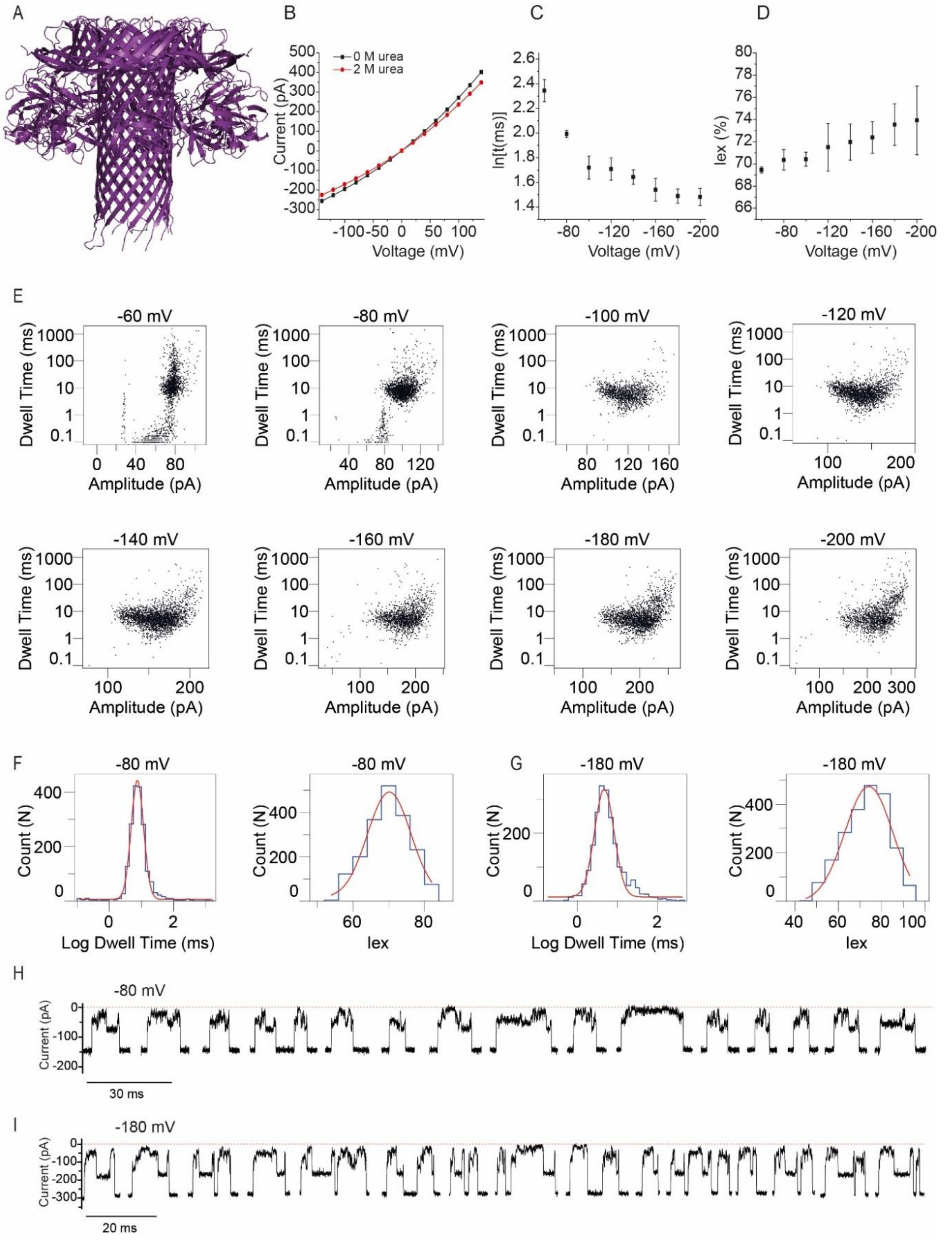

***Figure S5. Translocation malE219a through lysenin WT nanopore*.** ***A)*** *Cartoon representation of the lysenin nanopore.* ***B)*** *I-V curves in 1 M KCl at pH 7.5 of the lysenin nanopore in 0 M and 2 M urea. Each curve was obtained from a triplicate measurement.* ***C)*** *Dwell time dependence on the applied potential.* ***D)*** *$I_{ex\%}$ dependence on the applied potential. All data points represent the average of three independent experiments and the error bars corespond to standard deviation (SD).* ***E)*** *Scatter plots from -60 to -200 mV.* ***F-G)*** *Examples of histograms for the log(dwell time) and $I_{ex\%}$ at -80 mV (F) and -180 mV (G).* ***H-I)*** *Typical events at -80 mV (H) and -180 mV, respectively (I). Recordings were performed in 1 M KCl, 2 M urea, 15 mM HEPES, pH 7.5, 50 kHz sampling, 10 kHz Bessel filter. Each measurement was performed in a least a triplicate.*

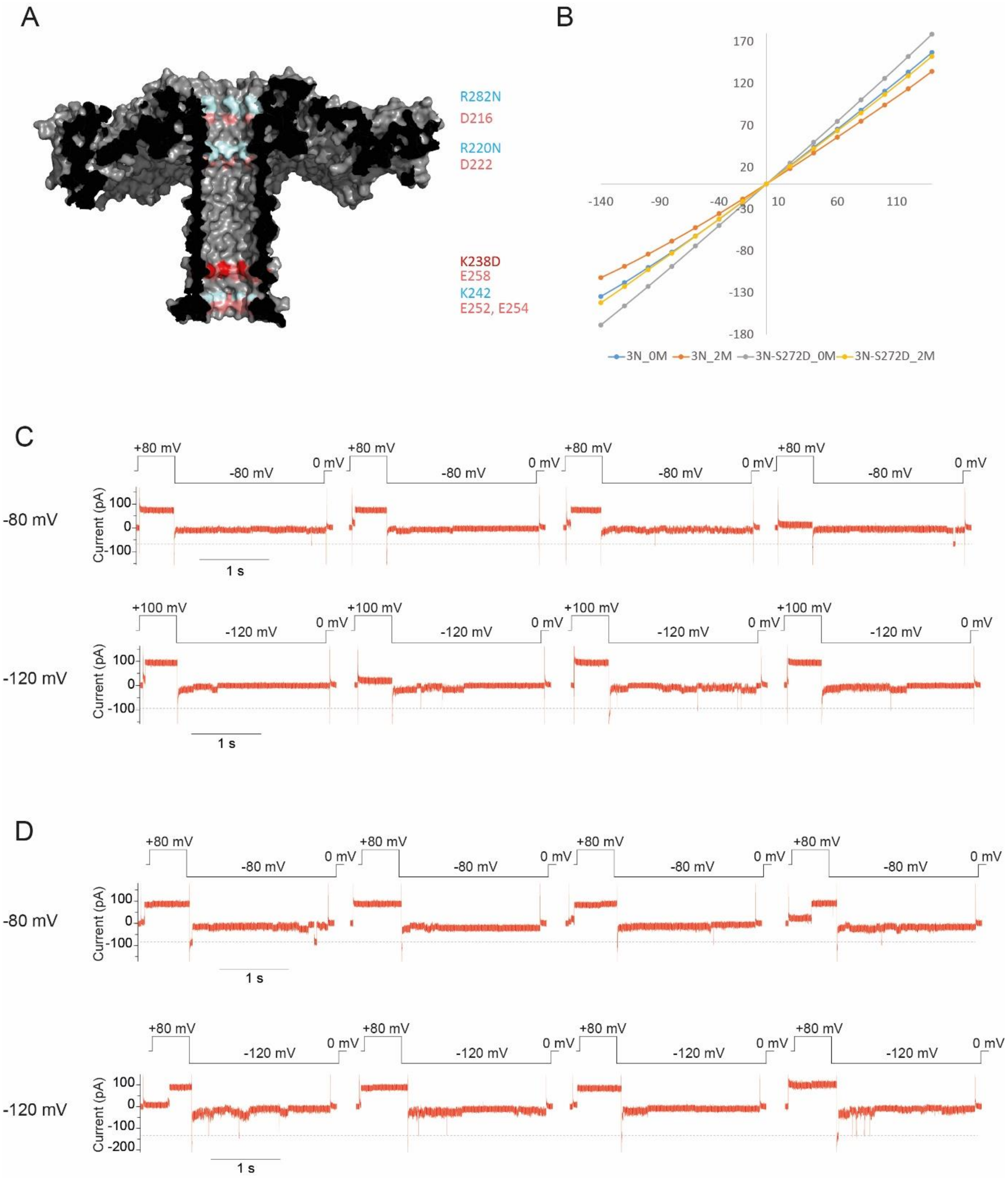

***Figure S6. Addition of malE219a to Aerolysin mutant nanopores in*** **cis in 2 M urea.** ***A****) Surface representation of the aerolysin nanopore, with positions of interest highlighted: R/K residues mutated to Asn are in blue, the Asp or Glu residues are in salmon, and K238D and S272D mutations are highlighted in red.* ***B****) I-V curves in 1 M KCl 15 mM HEPES pH 7.5 of the indicated mutants without and with 2 M urea (indicated as 0 M and 2M, respectively, in the figure inset).* ***C****) Aer 1D-3N at -80 and -100 mV.* ***D****) Aer 2D-3N at -80 mV and -120 mV. Recordings in panls C and D were performed in 1 M KCl, 2 M urea, 15 mM HEPES, pH 7.5, 50 kHz sampling, 10 kHz Bessel filter. Each measurement was performed in at least a triplicate.*

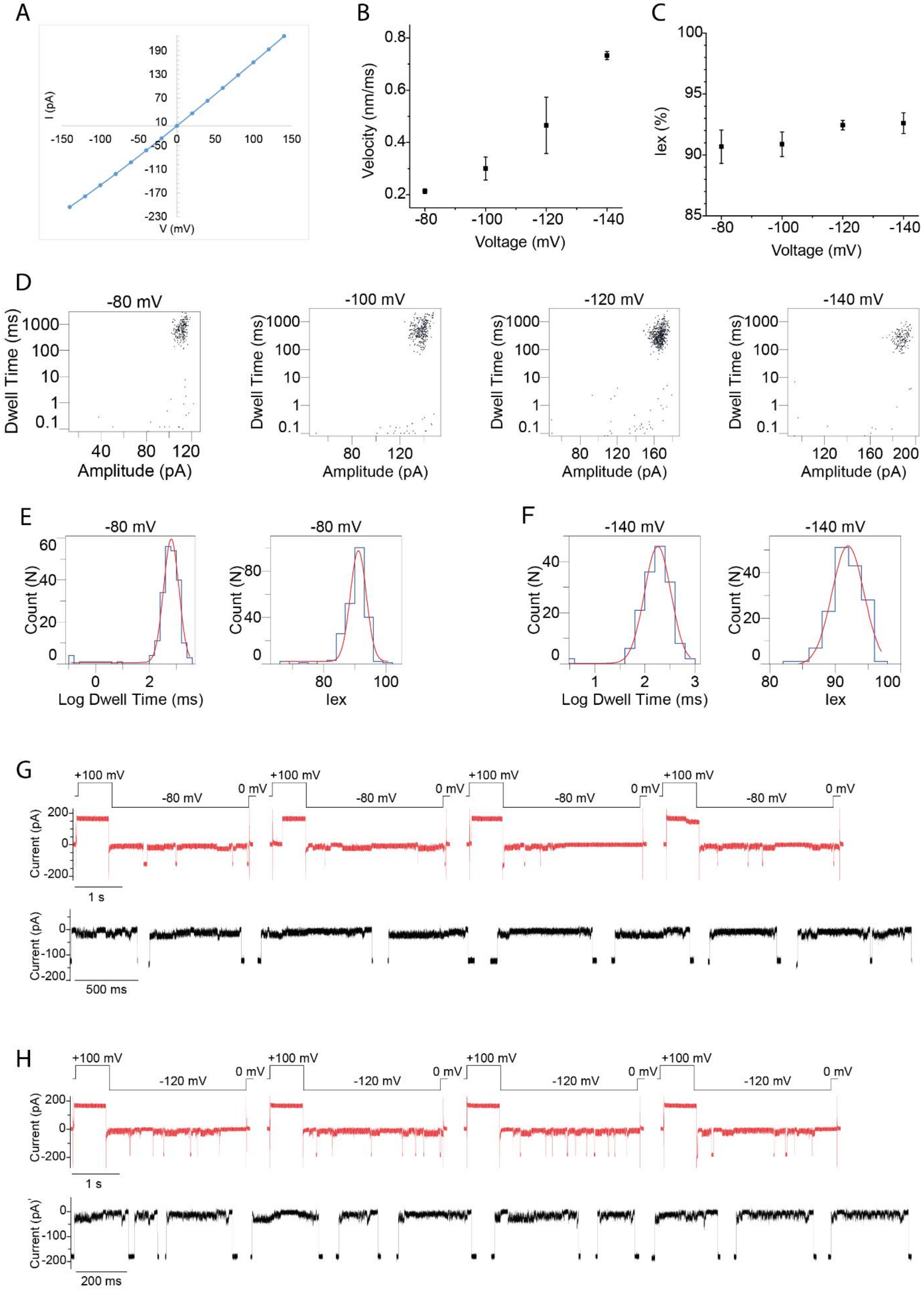

***Figure S7. Addition of malE219a to Aerolysin 1D-3N in cis in 1.8 M KCl. A****) I-V curve of the aerolysin 1D-3N mutant in 1.8 M KCl, 2 M urea, pH 7.5.* ***B****) Velocity dependence on the applied potential.* ***C)*** *$I_{ex\%}$ dependence on the applied potential. Data points represent averages from three independent experiments and the error bars correspond to standard deviations (SD).* ***D)*** *Scatter plots (dwell time vs amplitude) associated with the malE21a9 translocation*

*at the sampled potentials.* ***E-F)*** *Examples of histograms obtained for the log(dwell time) and $I_{ex\%}$ at -80 mV and -140 mV, respectively.* ***G-H)*** *Typical sweeps (applied potential protocol at the top) and translocation events at -80 mV (G) and -120 mV (H). Recordings were carried out in 1.8 M KCl, 15 mM HEPES, 2 M urea, pH 7.5, 50 kHz sampling and 10 kHz Bessel filter.*

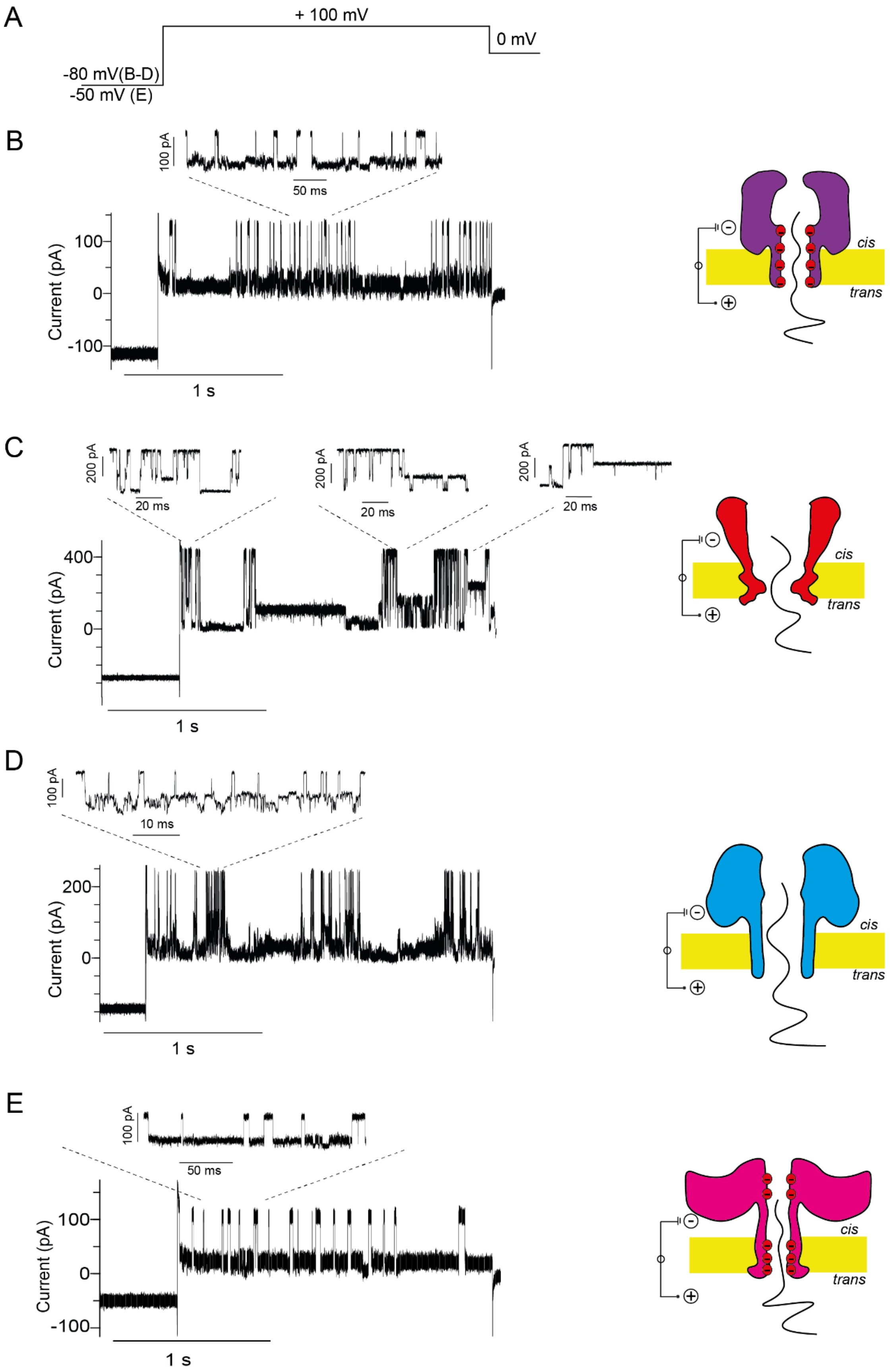

***Figure S8 Interaction of malE219a with nanopores from the trans*** **side.** ***A****) Applied potential in the sweeps protocol.* ***B****) CytK-4D nanopore.* ***C****) MspA WT nanopore.* ***D****) Lysenin WT nanopore.* ***E****) Aerolysin 1D-3N nanopore. The substrate was added to the* trans side.

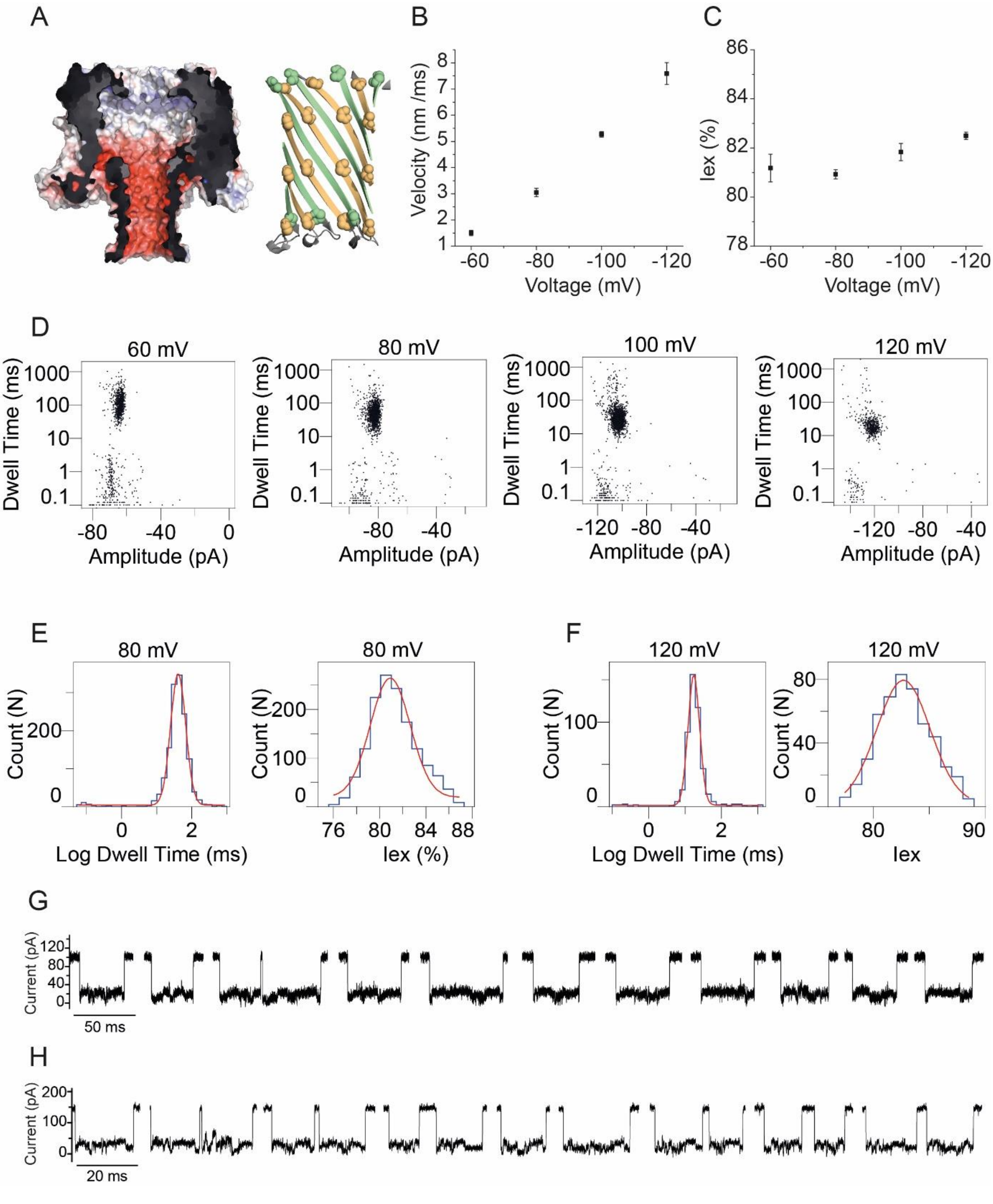

***Figure S9. Translocation malE219a from*** **trans to** **cis** ***through the CytK-4D nanopore. A)*** *Electrostatic map of the CytK nanopore at pH 7.5 in 1 M KCl (left) and the cartoon representation of the barrel where the substitutions with respect to the WT nanopore are highlighted (right).* ***B)*** *Velocity dependence on the applied potential.* ***C)*** *$I_{ex\%}$ dependence on the applied potential. Data points represent averages from three independent experiments and the error bars correspond to standard deviations (SD).* ***D)*** *Scatter plots (dwell time vs amplitude) associated with the malE219a translocation at the sampled potentials.* ***E-F)*** *Examples of histograms obtained for the log(dwell time) and $I_{ex\%}$ at +80 mV and +120 mV, respectively.* ***G-H)*** *Typical malE219a translocation events at +80 mV (G) and +120 mV (H). Recordings were carried out in 1 M KCl, 15 mM HEPES, 2 M urea, pH 7.5, 50 kHz sampling and 10 kHz Bessel filter and unlike other experiments, the substrate was added in* trans.

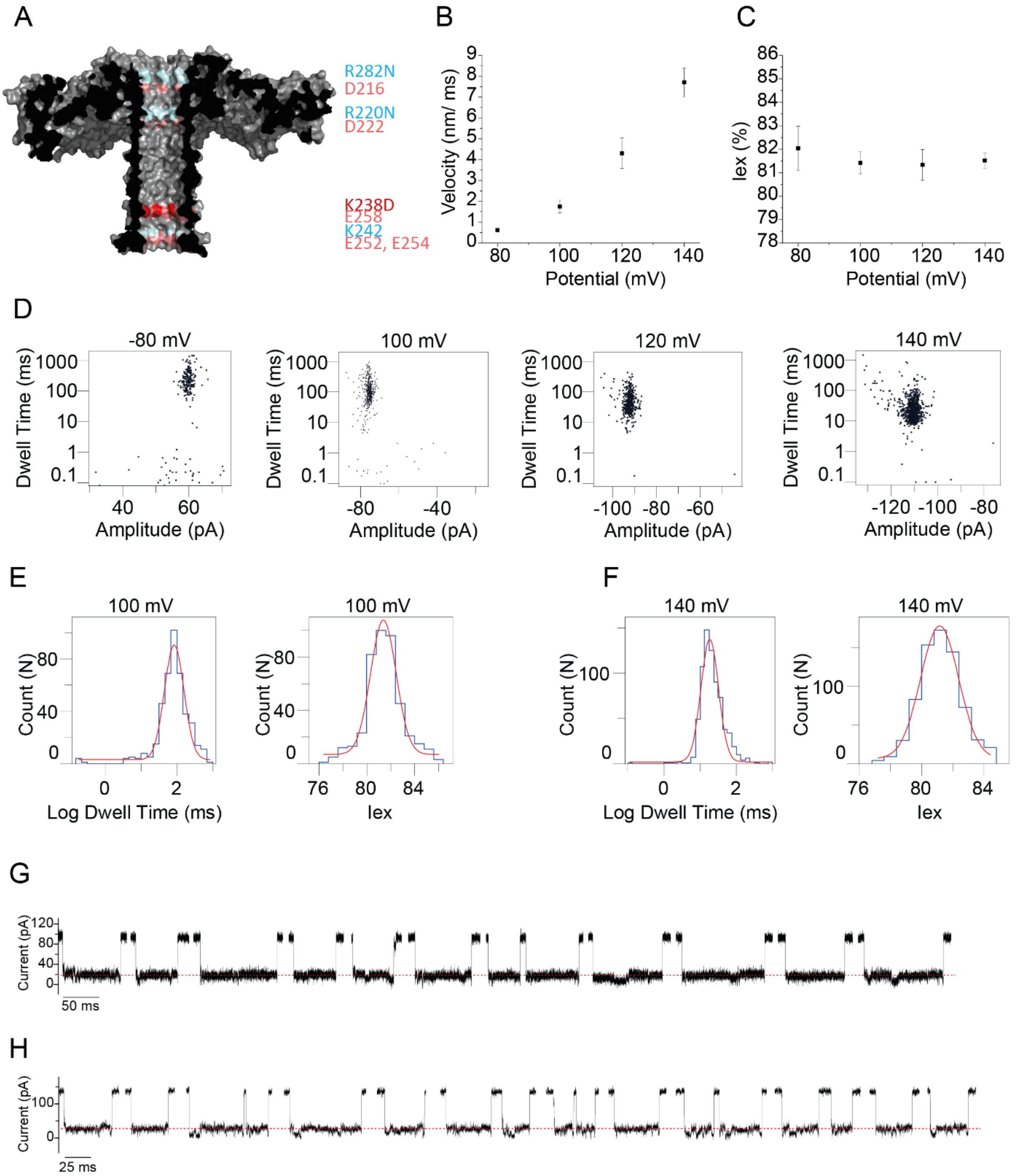

***Figure S10. Translocation malE219a from* trans *to* cis *through the Aer-1D-3N nanopore. A)*** *Surface representation of the aerolysin nanopore, with positions of interest highlighted: R/K mutated to N in blue, already existent Asp or Glu (salmon) and K238D (red).* ***B)*** *Velocity dependence on the applied potential.* ***C)*** *$I_{ex\%}$ dependence on the applied potential. Data points represent averages from three independent experiments and the error bars correspond to standard deviations (SD).* ***D)*** *Scatter plots (dwell time vs amplitude) associated with the malE219a translocation at the sampled potentials.* ***E-F)*** *Examples of histograms obtained for the log(dwell time) and $I_{ex}$ at +100 mV and +140 mV, respectively.* ***G-H)*** *Typical malE219a translocation events at +100 mV (H) and +140 mV (I). Recordings were carried out in 1 M KCl, 15 mM HEPES, 2 M urea, pH 7.5, 50 kHz sampling and 10 kHz Bessel filter and unlike other experiments, the substrate was added in* trans.

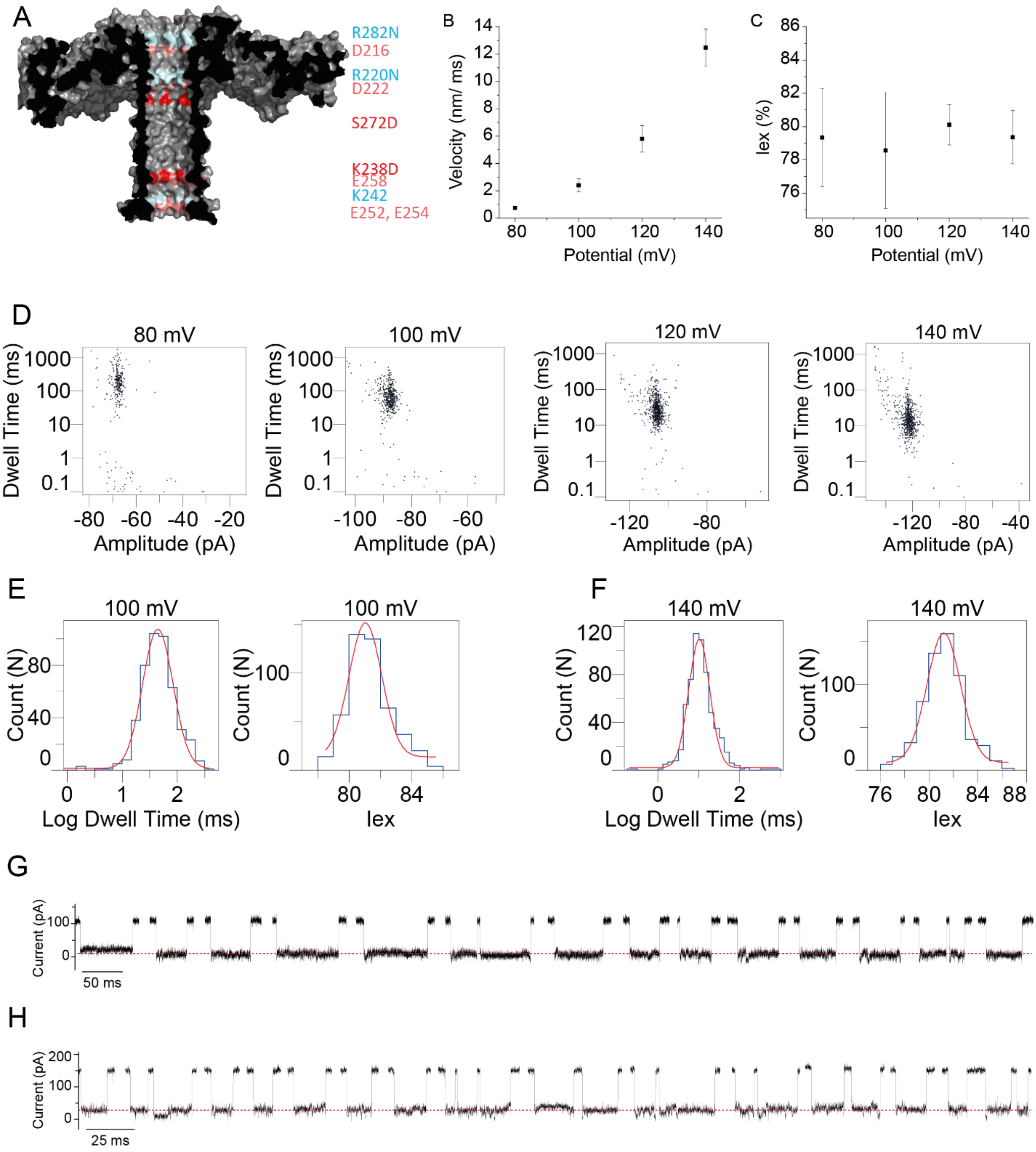

***Figure S11. Translocation malE219a from*** trans ***to*** cis ***through the Aer-2D-3N nanopore. A)*** *Surface representation of the aerolysin nanopore, with positions of interest highlighted: R/K mutated to N in blue, already existent Asp or Glu (salmon) and K238D and S272D (red).* ***B)*** *Velocity dependence on the applied potential.* ***C)*** *$I_{ex\%}$ dependence on the applied potential. Data points represent averages from three independent experiments and the error bars correspond to standard deviations (SD).* ***D)*** *Scatter plots (dwell time vs amplitude) associated with the malE219a translocation at the sampled potentials.* ***E-F)*** *Examples of histograms obtained for the log(dwell time) and $I_{ex\%}$ at 100 mV and 140 mV, respectively.* ***G-H)*** *Typical malE219a translocation events at +100 mV (**H**) and +140 mV (**I**). Recordings were carried out in 1 M KCl, 15 mM HEPES, 2 M urea, pH 7.5, 50 kHz sampling and 10 kHz Bessel filter and unlike other experiments, the substrate was added in* trans.

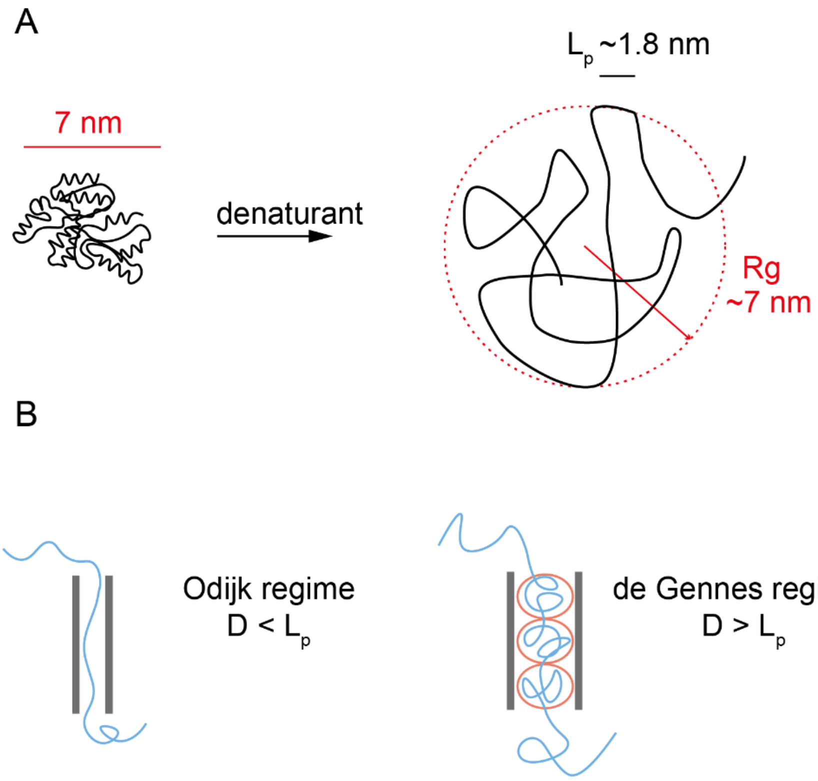

***Figure S12. Unstructured polymers in solution and in confinement. A****) In dilute protein solutions, urea induces the unfolding of malE219a, which then becomes an unstructured polymer described by a persistance length ($L_p$) and a gyration radius (Rg) as shown.* ***B****) The confinement of polymers inside nanochannels (and that of polypeptide through nanopores) can follow an Odijk regime if the nanochannel/nanopore diameter (D) is smaller than the persistence length of the polymer. If the pore D is much larger than the polymer $L_p$, confinement will follow a de Gennes regime in which case the polymer forms regular blobs (red circles) of unstructure polymer.*

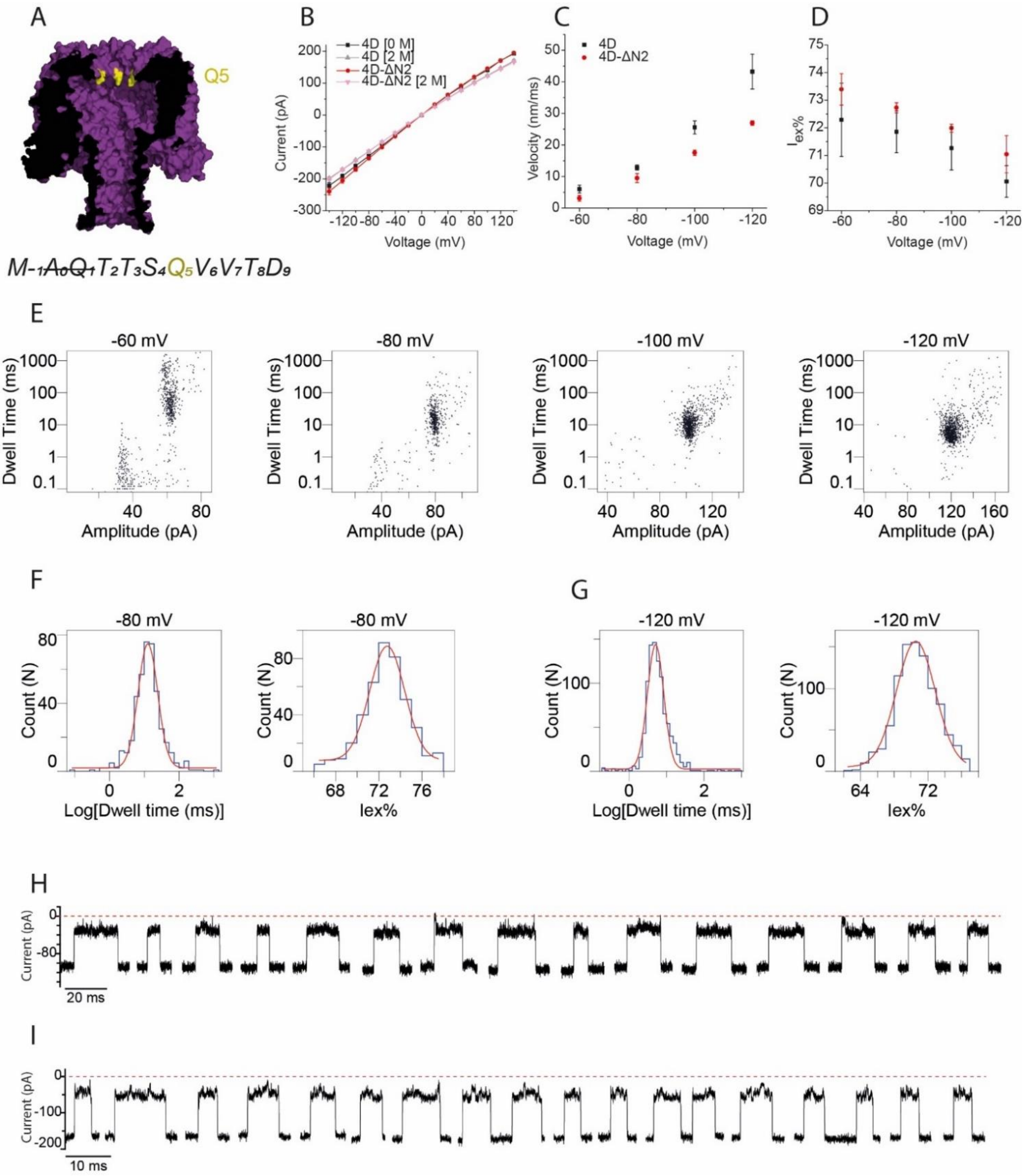

***Figure S13. Translocation of malE219a through the CytK-4D-ΔN2 nanopore. A)*** *The CytK nanopore, where the Q5 position is highlighted, accompanied by the sequence of the first N-term residues. In the case of this mutant* $A_0$ *and* $Q_1$ *are removed.* ***B)*** *IV curves in 1 M KCl at pH 7.5 of the 4D and 4D-ΔN2 mutants in 0 and 2 M urea; each curve was obtained from a triplicate measurement.* ***C)*** *Translocation velocity dependence of malE219a through the 4D and 4D-ΔN2 nanopores.* ***D)*** $I_{ex\%}$ *dependence of malE219a translocation through the 4D and 4D-ΔN2. Data points represent averages from three independent experiments and the error bars correspond to standard deviations (SD).* ***E)*** *Scatter plots (dwell time vs amplitude) associated with the malE219a translocation at the sampled potentials.* ***F-G)*** *Examples of histograms obtained for the log(dwell time) and* $I_{ex\%}$ *at -80 mV and -120 mV, respectively.* ***H-I)*** *Typical malE219a translocation events at -80 mV (**H**) and -120 mV (**I**). Recordings were carried out in 1 M KCl, 15 mM HEPES, 2 M urea, pH 7.5, 50 kHz sampling and 10 kHz Bessel filter.*

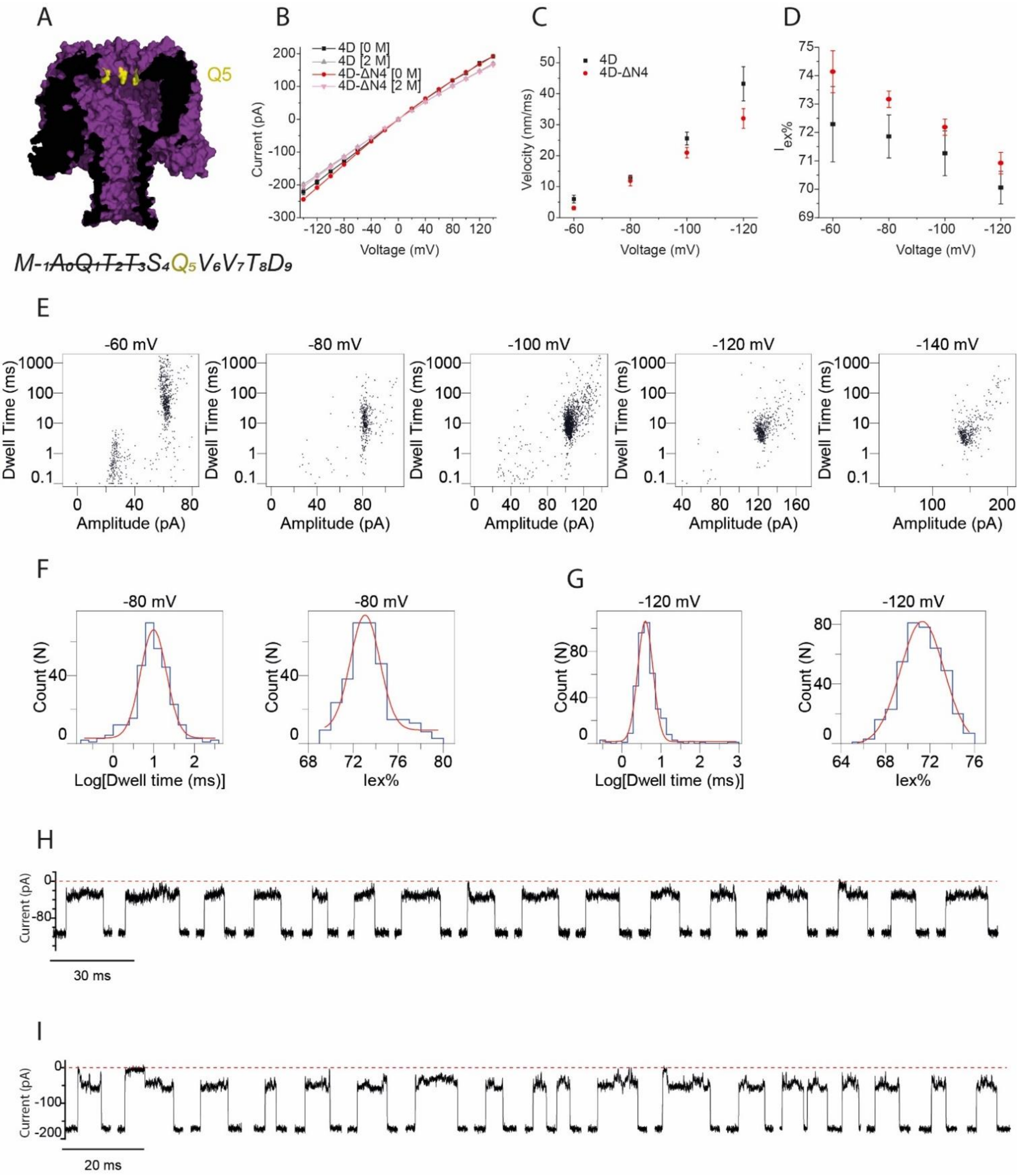

***Figure S14. Translocation of malE219a through the CytK-4D-ΔN4 nanopore. A)*** *The CytK nanopore, where the Q5 position is highlighted, accompanied by the sequence of the first N-term residues. In the case of this mutant* $A_0$ *–*$T_3$ *are removed.* ***B)*** *IV curves in 1 M KCl at pH 7.5 of the 4D and 4D-ΔN4 mutants in 0 and 2 M urea; each curve was obtained from a triplicate measurement.* ***C)*** *Translocation velocity dependence of malE219a through the 4D and 4D-ΔN4.* ***D)*** $I_{ex\%}$ *dependence of malE219a translocation through the 4D and 4D-ΔN4. Data points represent averages from three independent experiments and the error bars correspond to standard deviations (SD).* ***E)*** *Scatter plots (dwell time vs amplitude) associated with the malE219a translocation at the sampled potentials.* ***F-G)*** *Examples of histograms obtained for the log(dwell time) and* $I_{ex\%}$ *at -80 mV and -120 mV, respectively.* ***H-I)*** *Typical malE219a translocation events at -80 mV (**H**) and -120 mV (**I**). Recordings were carried out in 1 M KCl, 15 mM HEPES, 2 M urea, pH 7.5, 50 kHz sampling and 10 kHz Bessel filter.*

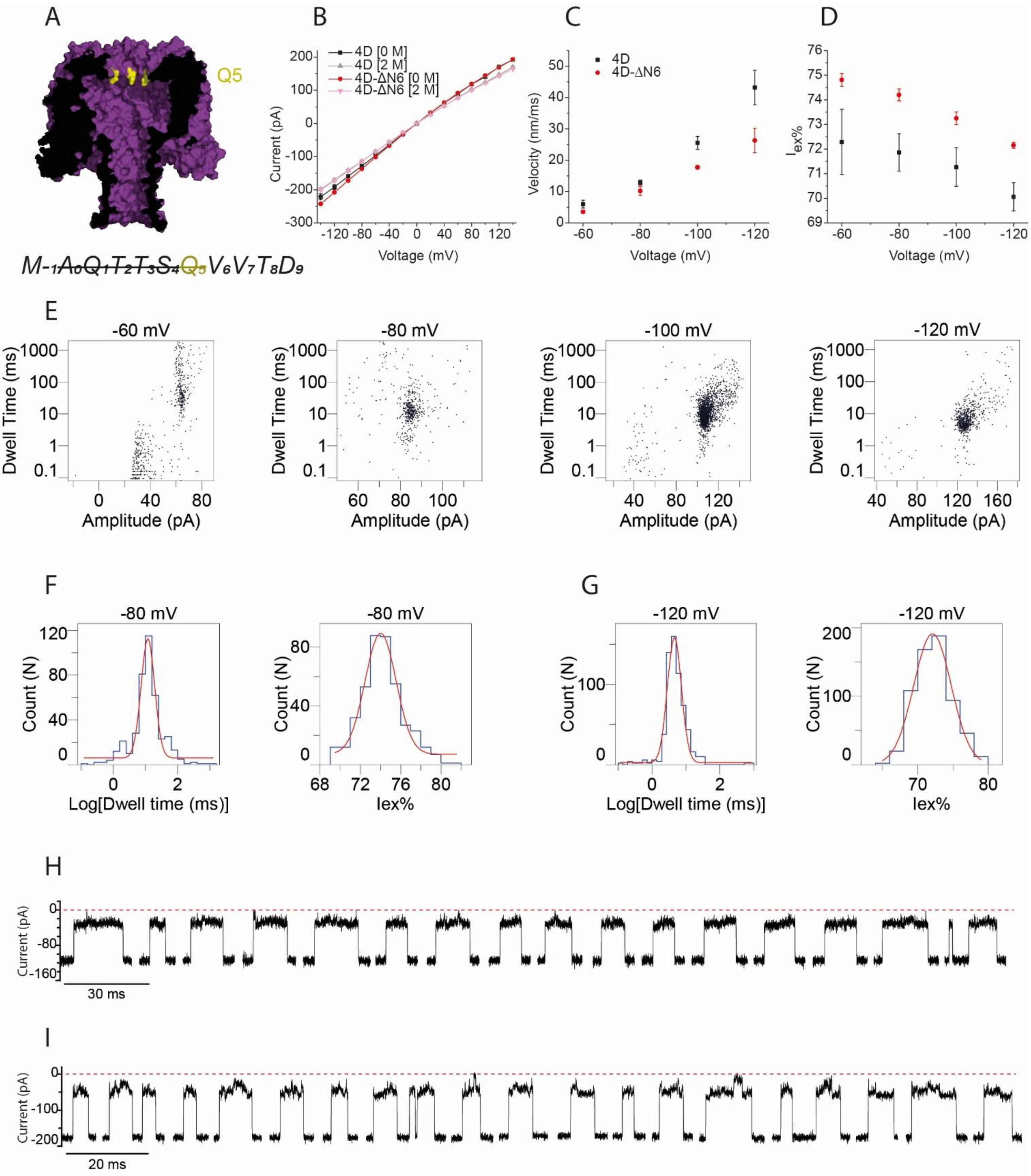

***Figure S15. Translocation of malE219a through the CytK-4D-ΔN6 nanopore. A)*** *The CytK nanopore, where the Q5 position is highlighted, accompanied by the sequence of the first N-term residues. In the case of this mutant* $A_0$ *–*$Q_5$ *are removed.* ***B)*** *IV curves in 1 M KCl at pH 7.5 of the 4D and 4D-ΔN6 mutants in 0 and 2 M urea; each curve was obtained from a triplicate measurement.* ***C)*** *Translocation velocity dependence of malE219a through the 4D and 4D-ΔN6.* ***D)*** $I_{ex\%}$ *dependence of malE219a translocation through the 4D and 4D-ΔN6. Data points represent averages from three independent experiments and the error bars correspond to standard deviations (SD).* ***E)*** *Scatter plots (dwell time vs amplitude) associated with the malE219a translocation at the sampled potentials.* ***F-G)*** *Examples of histograms obtained for the log(dwell time) and* $I_{ex\%}$ *at -80 mV and -120 mV, respectively.* ***H-I)*** *Typical malE219a translocation events at -80 mV (H) and -120 mV (I). Recordings were carried out in 1 M KCl, 15 mM HEPES, 2 M urea, pH 7.5, 50 kHz sampling and 10 kHz Bessel filter.*

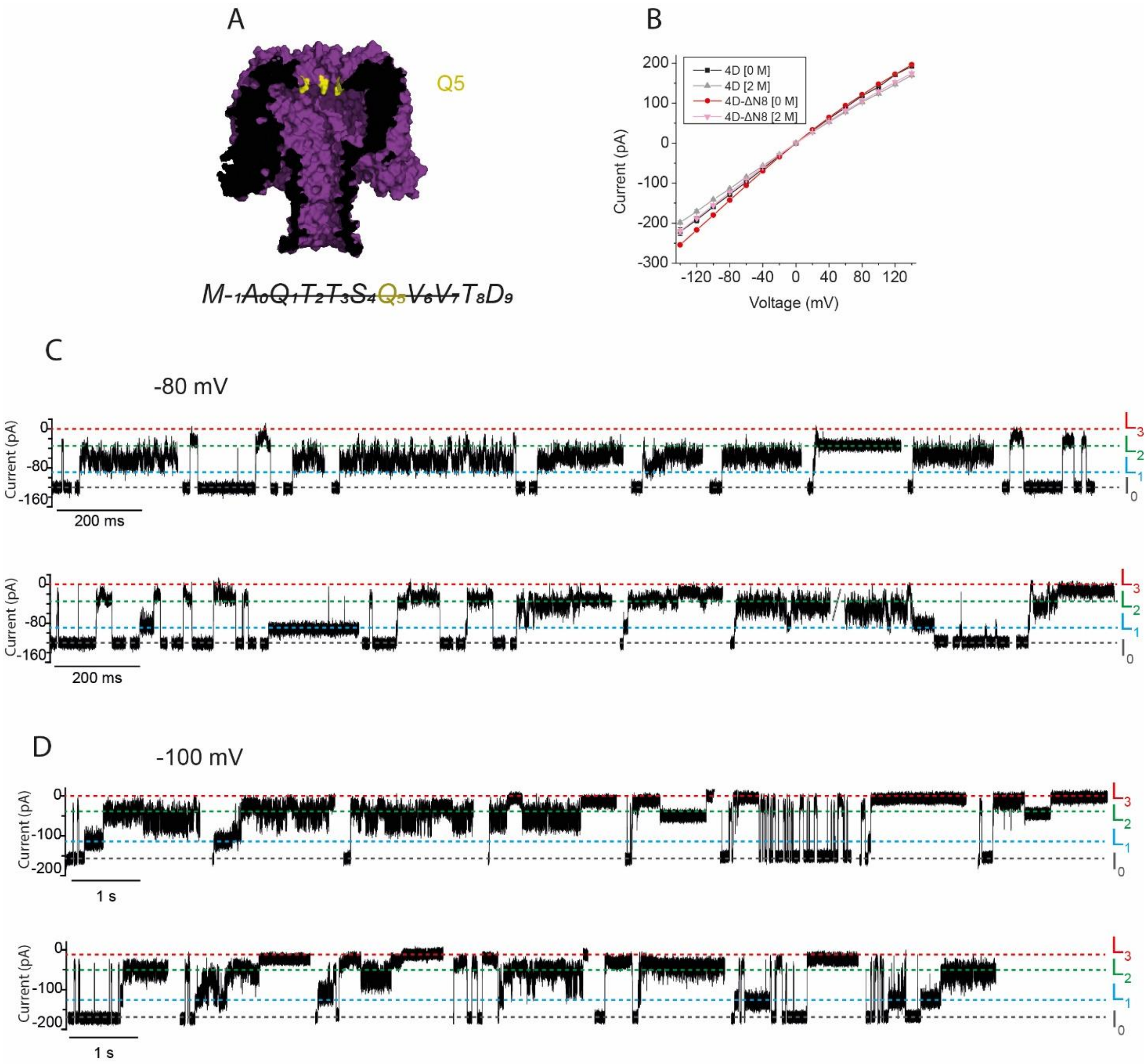

***Figure S16. Translocation of malE219a through the CytK-4D-ΔN8 nanopore. A)*** *The CytK nanopore, where the Q5 position is highlighted, accompanied by the sequence of the first N-term residues. In the case of this mutant $A_0$-$V_7$ are removed, which causes the widening of the pore entry.* ***B)*** *IV curves in 1 M KCl at pH 7.5 of the 4D and 4D-ΔN8 mutants in 0 and 2 M urea; each curve was obtained from a triplicate measurement.* ***C-D)*** *Typical malE219a translocation events at -80 m (**C**) and -100 mV (**D**). Recordings were carried out in 1 M KCl, 15 mM HEPES, 2 M urea, pH 7.5, 50 kHz sampling rate and 10 kHz Bessel filter. Translocation events (level marked with green) are less predominant in the case of this mutant; instead, long-lived events of various levels are enriched. Occasionally, at low potential, shallow capture events, which are also spontaneously released, occur; these events may be attributed to the substrate coiling at the entry and transiently blocking it, followed by its release, instead of progression through the vestibule. The more prominent capture events start with a shallow level (blue) and are followed by a deeper, noisier level (between levels marked with blue and green at -80 mV, or around the green level at -100 mV) and are relatively long-lived (>1-2 s).*

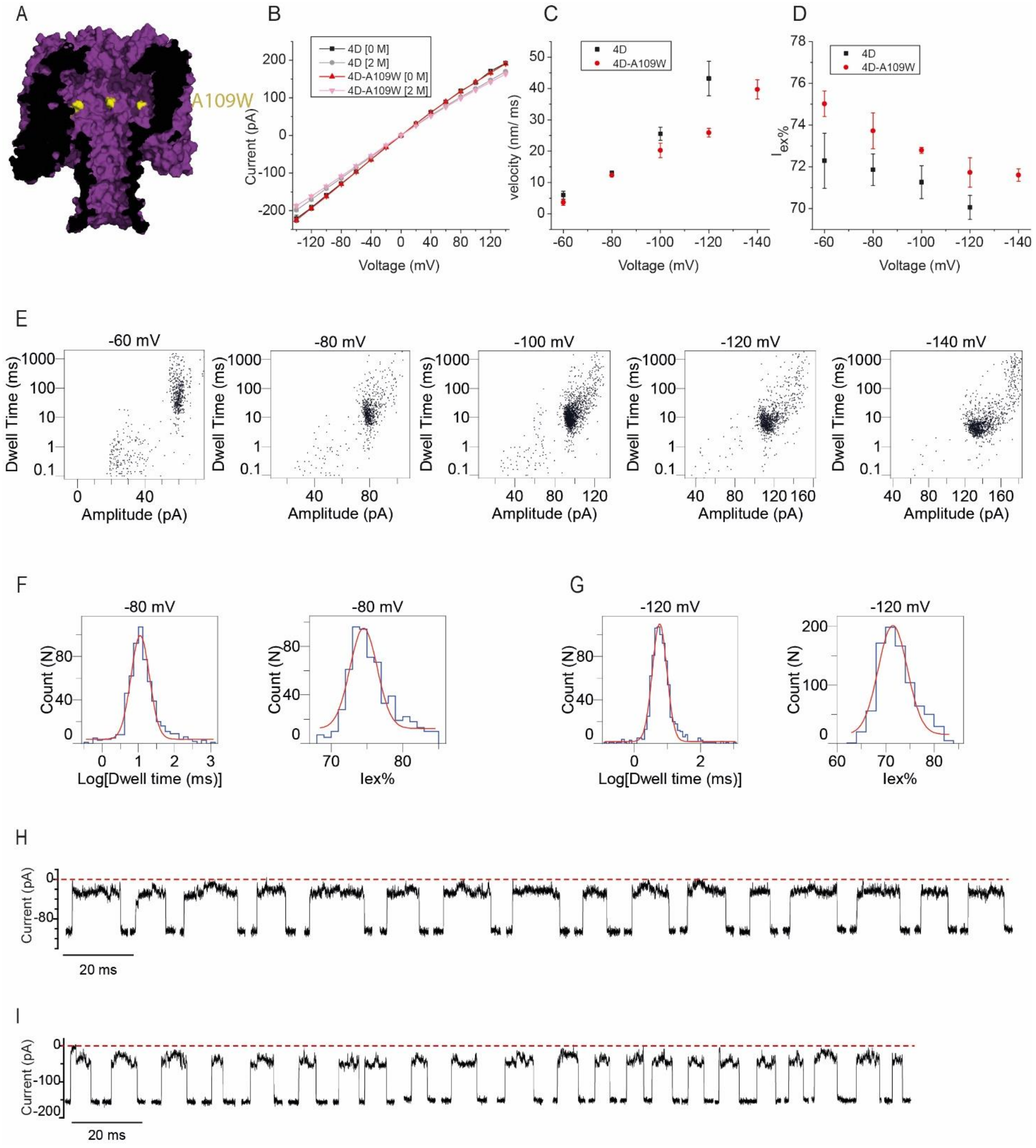

***Figure S17**. **Translocation of malE219a through the CytK-4D-A109W nanopore. A)** The CytK nanopore, where the A109W position is highlighted. **B)** IV curves in 1 M KCl at pH 7.5 of the 4D and 4D-A109W mutants in 0 and 2 M urea; each curve was obtained from a triplicate measurement. **C)** Translocation velocity dependence of malE219a through the 4D and 4D-A109W. **D)** $I_{ex\%}$ dependence of malE219a translocation through the 4D and 4D-A109W. Data points represent averages from three independent experiments and the error bars correspond to standard deviations (SD). **E)** Scatter plots (dwell time vs amplitude) associated with the malE219a translocation at the sampled potentials. **F-G)** Examples of histograms obtained for the log(dwell time) and $I_{ex\%}$ at -80 mV and -120 mV, respectively. **H-I)** Typical malE219a translocation events at -80 mV (**H**) and -120 mV (**I**). Recordings were carried out in 1 M KCl, 15 mM HEPES, 2 M urea, pH 7.5, 50 kHz sampling and 10 kHz Bessel filter.*

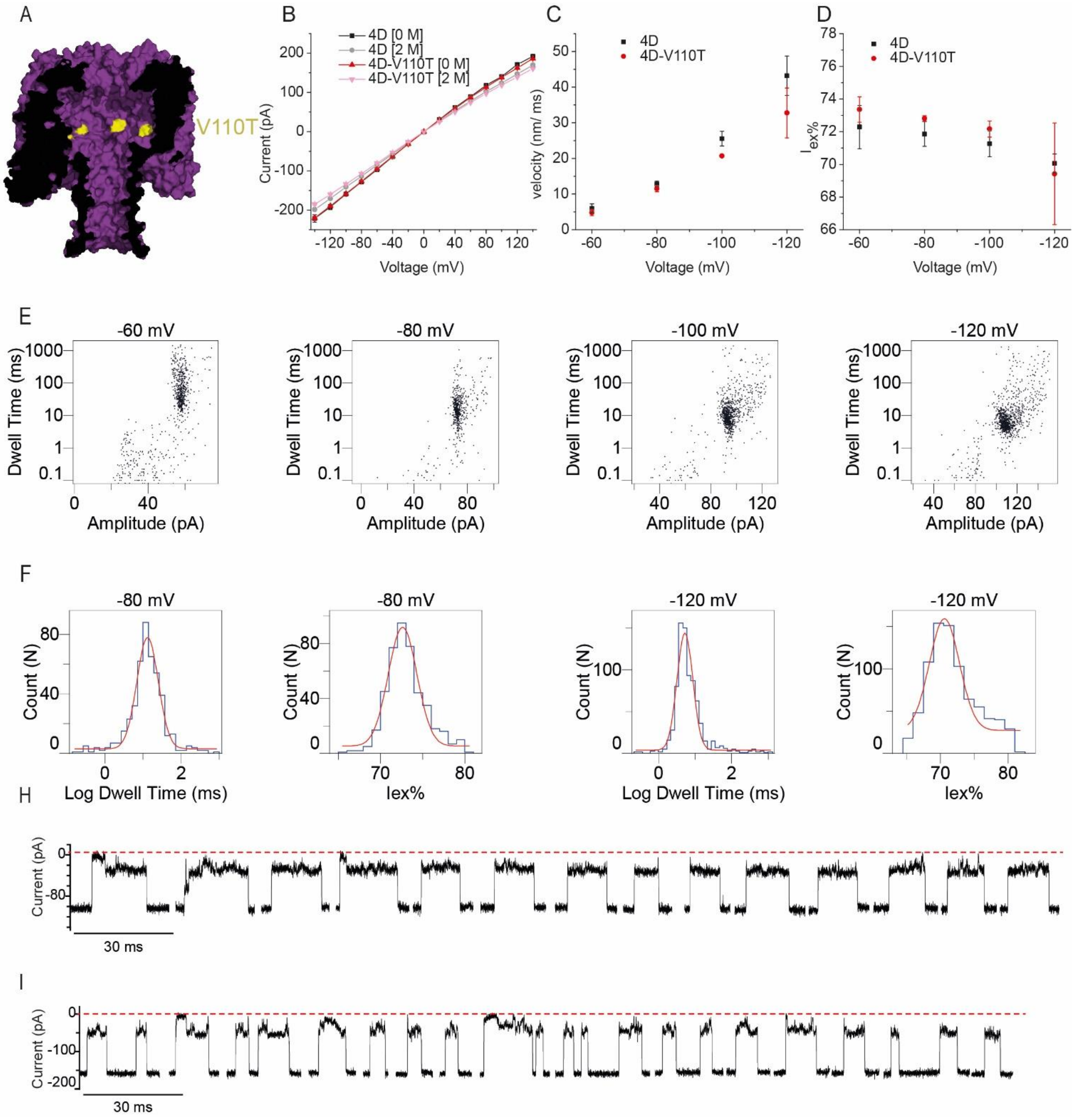

***Figure S18. Translocation of malE219a through the CytK-4D-V110T nanopore. A)*** *The CytK nanopore, where the V110T position is highlighted.* ***B)*** *IV curves in 1 M KCl at pH 7.5 of the 4D and 4D-A109W mutants in 0 and 2 M urea; each curve was obtained from a triplicate measurement.* ***C)*** *Translocation velocity dependence of malE219a through the 4D and 4D-V110T.* ***D)*** *$I_{ex\%}$ dependence of malE219a translocation through the 4D and 4D-V110T. Data points represent averages from three independent experiments and the error bars correspond to standard deviations (SD).* ***E)*** *Scatter plots (dwell time vs amplitude) associated with the malE219a translocation at the sampled potentials.* ***F-G)*** *Examples of histograms obtained for the log(dwell time) and $I_{ex\%}$ at -80 mV and -120 mV, respectively.* ***H-I)*** *Typical malE219a translocation events at -80 mV (**H**) and -120 mV (**I**). Recordings were carried out in 1 M KCl, 15 mM HEPES, 2 M urea, pH 7.5, 50 kHz sampling and 10 kHz Bessel filter.*

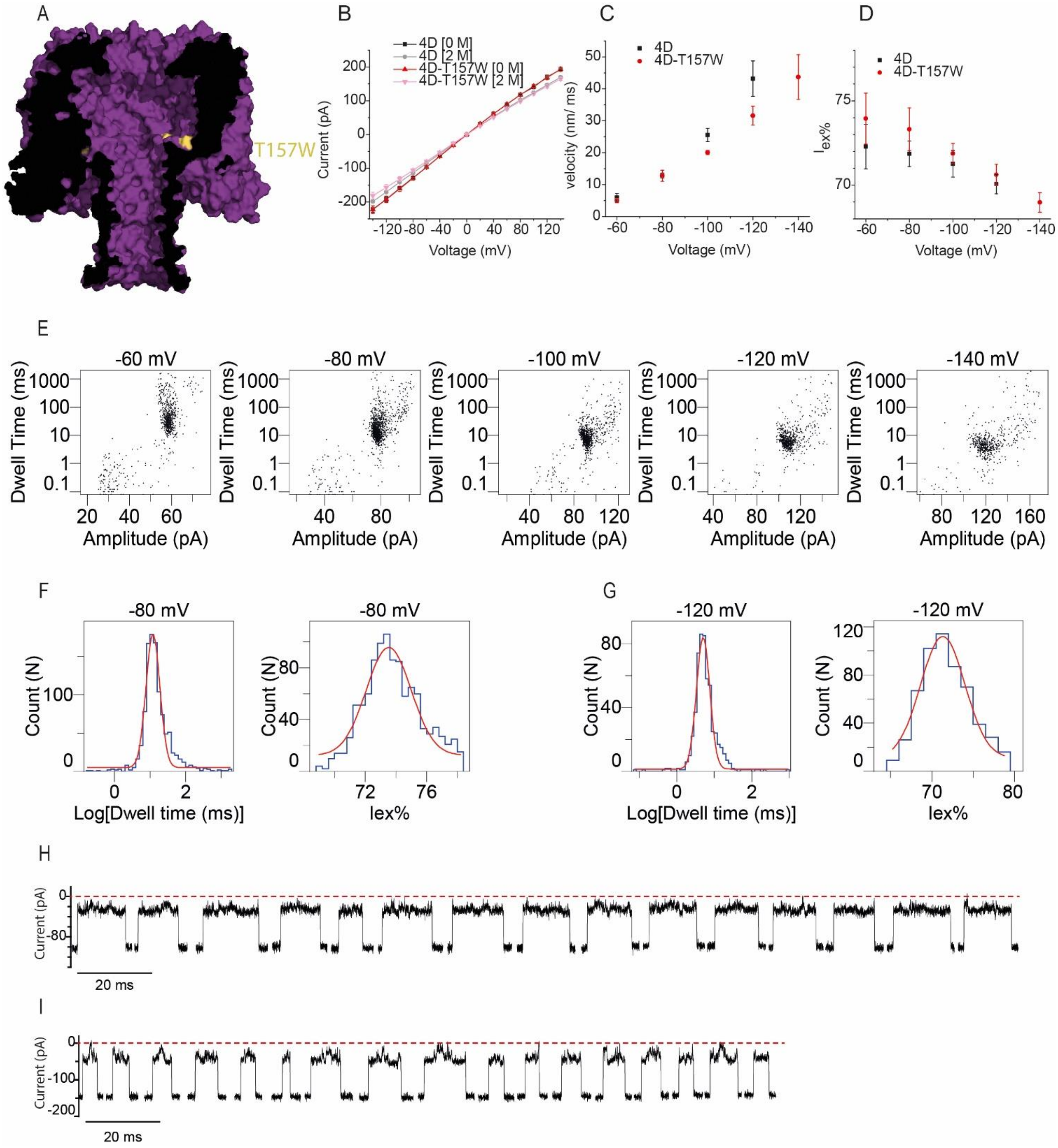

***Figure S19. Translocation of malE219a through the CytK-4D-T157W nanopore. A)*** *The CytK nanopore, where the T157W position is highlighted.* ***B)*** *IV curves in 1 M KCl at pH 7.5 of the 4D and 4D-T157W mutants in 0 and 2 M urea; each curve was obtained from a triplicate measurement.* ***C)*** *Translocation velocity dependence of malE219a through the 4D and 4D-T157W.* ***D)*** *$I_{ex\%}$ dependence of malE219a translocation through the 4D and 4D-T157W. Data points represent averages from three independent experiments and the error bars correspond to standard deviations (SD).* ***E)*** *Scatter plots (dwell time vs amplitude) associated with the malE219a translocation at the sampled potentials.* ***F-G)*** *Examples of histograms obtained for the log(dwell time) and $I_{ex\%}$ at -80 mV and -120 mV, respectively.* ***H-I)*** *Typical malE219a translocation events at -80 mV (**H**) and -120 mV (**I**). Recordings were carried out in 1 M KCl, 15 mM HEPES, 2 M urea, pH 7.5, 50 kHz sampling and 10 kHz Bessel filter.*

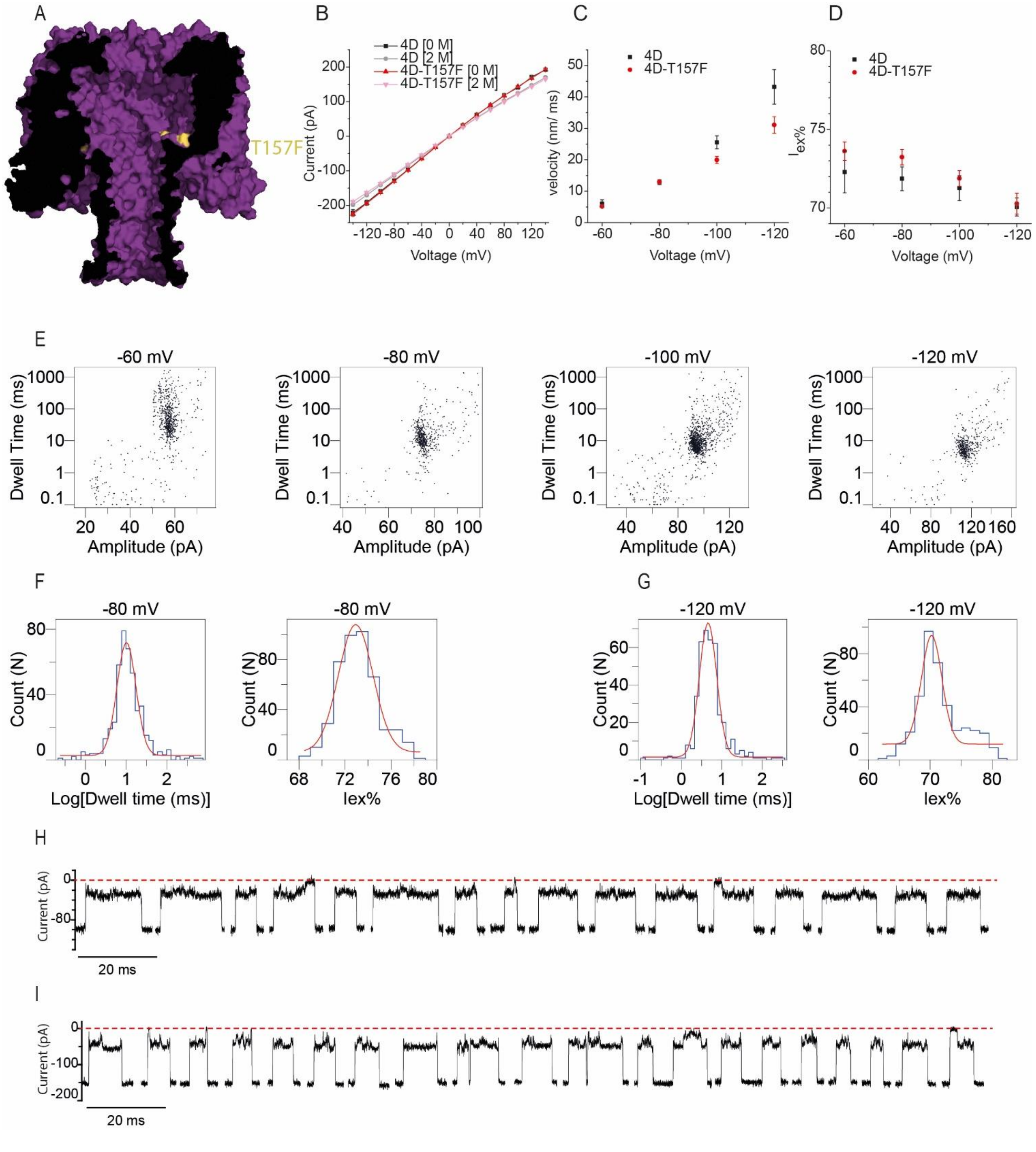

***Figure S20. Translocation of malE219a through the CytK-4D-T157F nanopore. A)*** *The CytK nanopore, where the T157F position is highlighted.* ***B)*** *IV curves in 1 M KCl at pH 7.5 of the 4D and 4D-T157F mutants in 0 and 2 M urea; each curve was obtained from a triplicate measurement.* ***C)*** *Translocation velocity dependence of malE219a through the 4D and 4D-T157F.* ***D)*** *$I_{ex\%}$ dependence of malE219a translocation through the 4D and 4D-T157F. Data points represent averages from three independent experiments and the error bars correspond to standard deviations (SD).* ***E)*** *Scatter plots (dwell time vs amplitude) associated with the malE219a translocation at the sampled potentials.* ***F-G)*** *Examples of histograms obtained for the log(dwell time) and $I_{ex\%}$ at -80 mV and -120 mV, respectively.* ***H-I)*** *Typical malE219a translocation events at -80 mV (**H**) and -120 mV (**I**). Recordings were carried out in 1 M KCl, 15 mM HEPES, 2 M urea, pH 7.5, 50 kHz sampling and 10 kHz Bessel filter.*

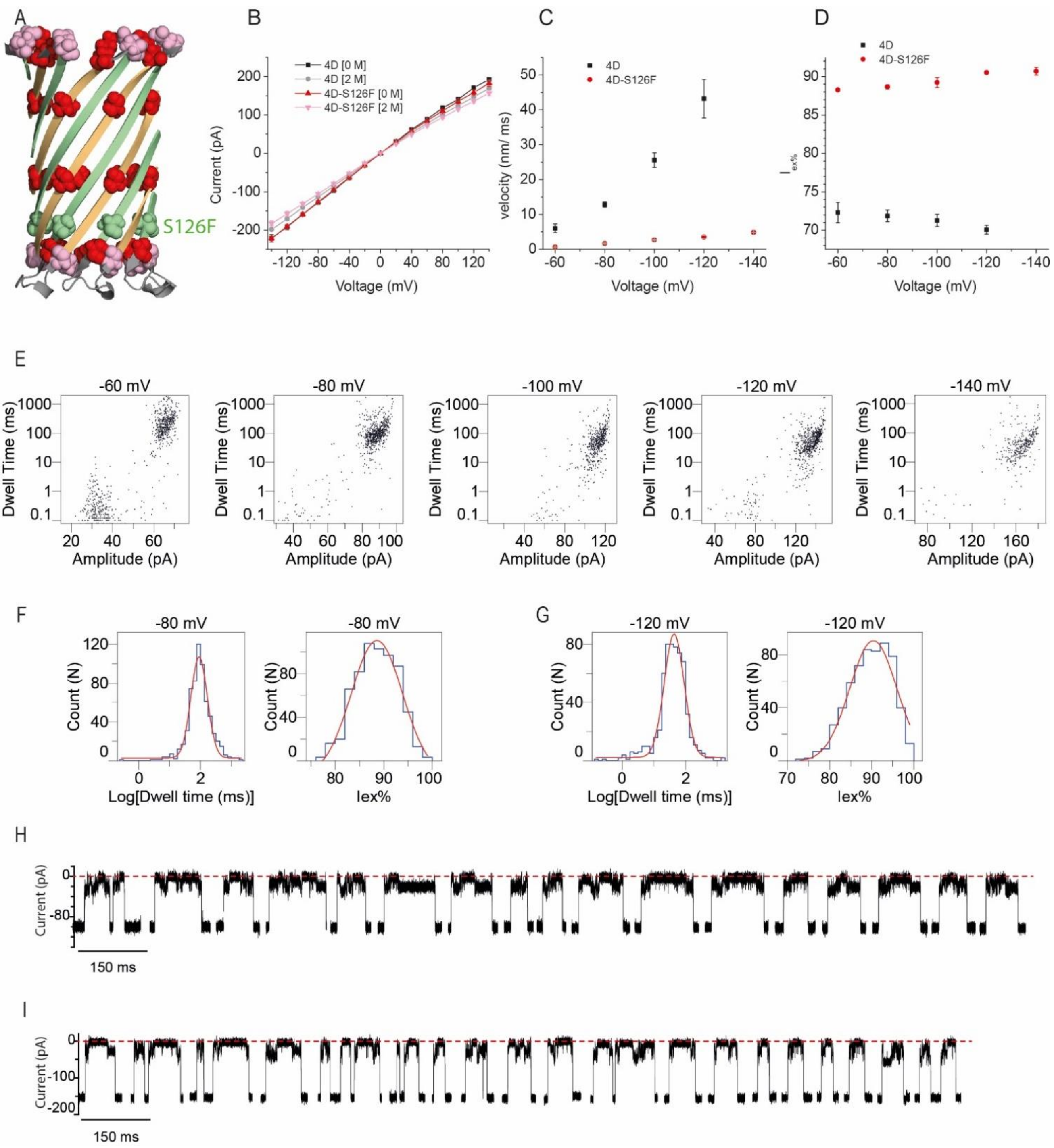

***Figure S21. Translocation of malE219a through the CytK-4D-S126F nanopore. A)*** *The barrel of the CytK nanopore. The N-term strand is depicted in green, while the C-term strand in green. The relevant residues are shown as spheres: E112 and E139 in pink, the four aspartate substitutions in red (K128D, Q145D, S151D and K155D) and S126 in green.* ***B)*** *IV curves in 1 M KCl at pH 7.5 of the 4D and 4D-S126F mutants in 0 and 2 M urea; each curve was obtained from a triplicate measurement.* ***C)*** *Translocation velocity dependence of malE219a through the 4D and 4D-S126F.* ***D)*** $I_{ex\%}$ *dependence of malE219a translocation through the 4D and 4D-S126F. Data points represent averages from three independent experiments and the error bars correspond to standard deviations (SD).* ***E)*** *Scatter plots (dwell time vs amplitude) associated with the malE219a translocation at the sampled potentials.* ***F-G)*** *Examples of histograms obtained for the log(dwell time) and* $I_{ex\%}$ *at -80 mV and -120 mV, respectively.* ***H-I)*** *Typical malE219a translocation events at -80 mV (**H**) and -120 mV (**I**). Recordings were carried out in 1 M KCl, 15 mM HEPES, 2 M urea, pH 7.5, 50 kHz sampling and 10 kHz Bessel filter.*

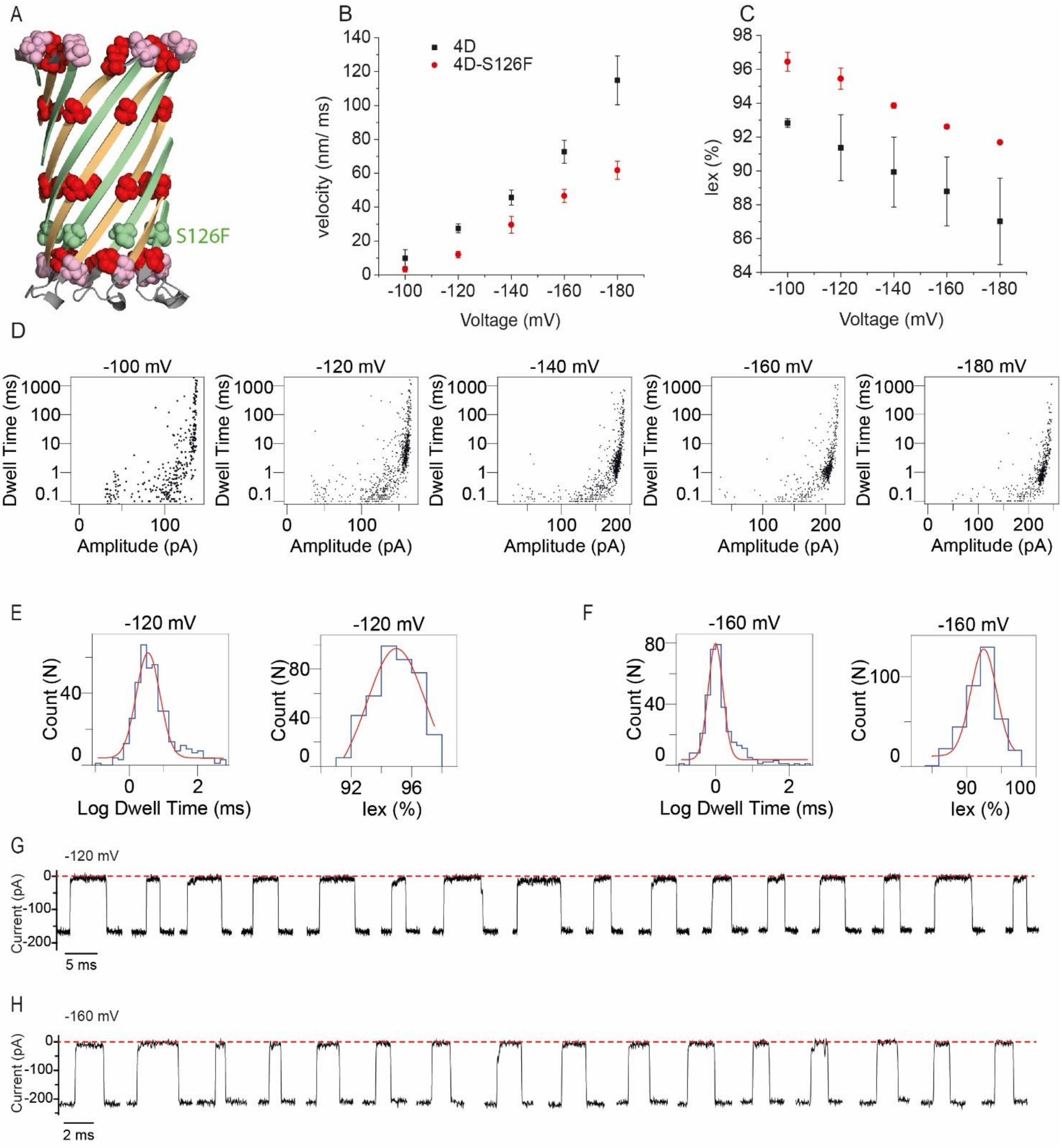

***Figure S22. Translocation of tzatziki through the CytK-4D-S126F nanopore. A)*** *The barrel of the CytK nanopore. The N-term strand is depicted in green, while the C-term strand in green. The relevant residues are shown as spheres: E112 and E139 in pink, the four Asp positions in red (K128D, Q145D, S151D and K155D) and S126 in green.* ***B)*** *Translocation velocity dependence of tzatziki through the 4D and 4D-S126F.* ***C)*** *$I_{ex\%}$ dependence of tzatziki translocation through the 4D and 4D-S126F. Data points represent averages from three independent experiments and the error bars correspond to standard deviations (SD).* ***D)*** *Scatter plots (dwell time vs amplitude) associated with the tzatziki translocation at the sampled potentials.* ***E-F)*** *Examples of histograms obtained for the log(dwell time) and $I_{ex\%}$ at -120 mV and -160 mV, respectively.* ***G-H)*** *Typical tzatziki translocation events at -120 mV (**G**) and -160 mV (**H**). Recordings were carried out in 1 M KCl, 15 mM HEPES, pH 7.5, 50 kHz sampling and 10 kHz Bessel filter.*

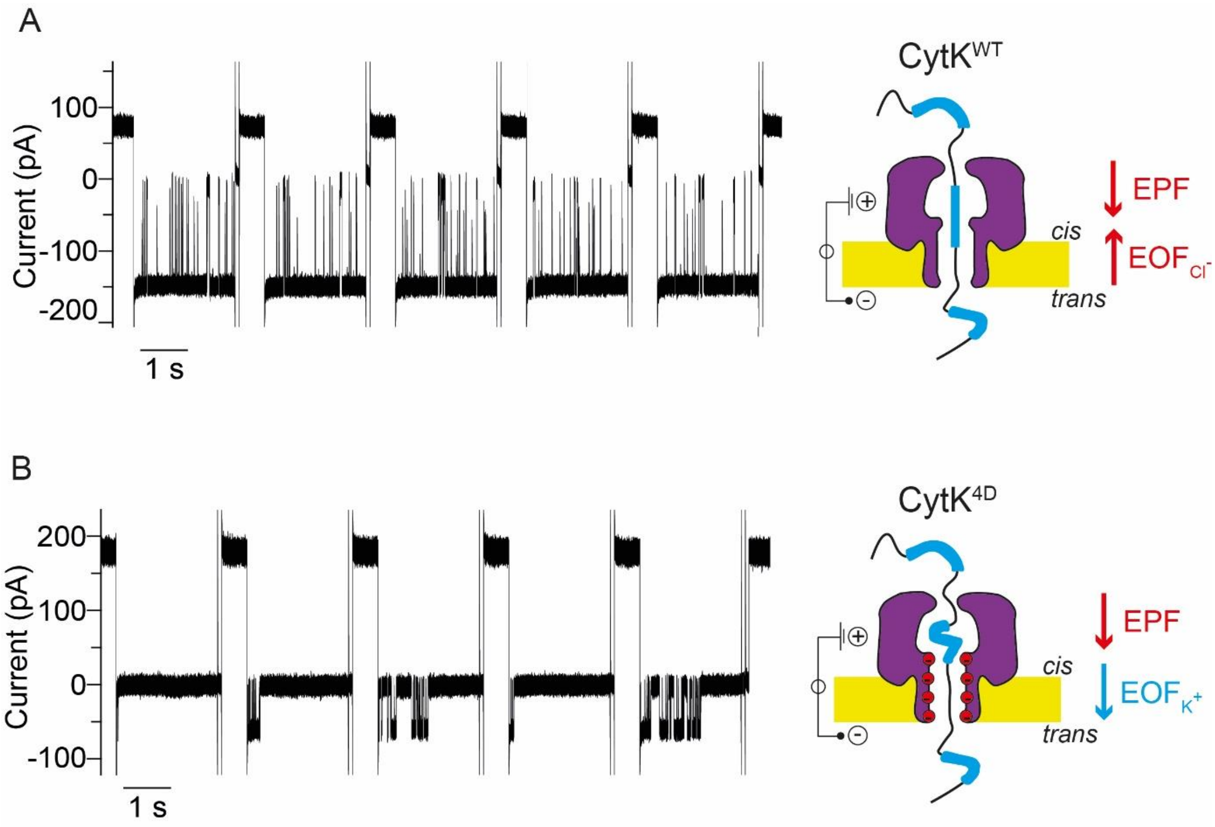

***Figure S23. EPF-driven versus EPF-EOF-driven transport of S1 across CytK nanopores.*** *The left panels show 5 sweeps for S1 translocation across CytK-WT (-160 mV, panel* ***A****) or CytK-4D (-40 mV, panel* ***B****) nanopores. Each sweep begins with a positive applied potential and continues with the negative applied potential. -160 mV (CytK-WT) and -40 mV (CytK-4D) were chosen because the dwell times are approximately the same, therefore the substrate experiences the same velocity as it translocates the two nanopores. In the case of the WT nanopore, the substrate is unlikely to coil within the vestibule due to EPF and EOF being of opposite directions and no prolonged events are detected. By contrast, when the EPF and EOF are not opposing, long-lived events are observed. The latter might result from the sticking of arginine residues in S1 to the aspartate rings in CytK-4D, and/or the substrate coiling by the absence of an EOF opposing the electrophoretic translocation of S1.*

**References**

1. Bitinaite, J. *et al.* USERTM friendly DNA engineering and cloning method by uracil excision. *Nucleic Acids Res* **35**, 1992–2002 (2007).

2. Cavaleiro, A. M., Kim, S. H., Seppälä, S., Nielsen, M. T. & Nørholm, M. H. H. Accurate DNA Assembly and Genome Engineering with Optimized Uracil Excision Cloning. *ACS Synth Biol* **4**, 1042–1046 (2015).

3. Nørholm, M. H. A mutant Pfu DNA polymerase designed for advanced uracil-excision DNA engineering. *BMC Biotechnol* **10**, 21 (2010).

4. Versloot, R. C. A., Straathof, S. A. P., Stouwie, G., Tadema, M. J. & Maglia, G. β-Barrel Nanopores with an Acidic–Aromatic Sensing Region Identify Proteinogenic Peptides at Low pH. *ACS Nano* **16**, 7258–7268 (2022).

5. Sauciuc, A., Morozzo della Rocca, B., Tadema, M. J., Chinappi, M. & Maglia, G. Translocation of linearized full-length proteins through an engineered nanopore under opposing electrophoretic force. *Nat Biotechnol* (2023) doi:10.1038/s41587-023-01954-x.

6. Kabsch, W. *XDS*. *Acta Crystallogr D Biol Crystallogr* **66**, 125–132 (2010).

7. McCoy, A. J. *et al. Phaser* crystallographic software. *J Appl Crystallogr* **40**, 658–674 (2007).

8. Emsley, P., Lohkamp, B., Scott, W. G. & Cowtan, K. Features and development of *Coot*. *Acta Crystallogr D Biol Crystallogr* **66**, 486–501 (2010).

9. Liebschner, D. *et al.* Macromolecular structure determination using X-rays, neutrons and electrons: recent developments in *Phenix*. *Acta Crystallogr D Struct Biol* **75**, 861–877 (2019).

10. Pettersen, E. F. *et al.* <scp>UCSF ChimeraX</scp> : Structure visualization for researchers, educators, and developers. *Protein Science* **30**, 70–82 (2021).

11. Maglia, G., Heron, A. J., Stoddart, D., Japrung, D. & Bayley, H. Analysis of Single Nucleic Acid Molecules with Protein Nanopores. in 591–623 (2010). doi:10.1016/S0076-6879(10)75022-9.
